# Designing and bioengineering of CDRs with higher affinity against receptor-binding domain (RBD) of SARS-CoV-2 Omicron variant

**DOI:** 10.1101/2024.09.19.613860

**Authors:** Vishakha Singh, Shweta Choudhary, Mandar Bhutkar, Sanketkumar Nehul, Sabika Ali, Jitin Singla, Pravindra Kumar, Shailly Tomar

## Abstract

The emergence of the SARS-CoV-2 Omicron variant highlights the need for innovative strategies to address evolving viral threats. This study bioengineered three nanobodies H11-H4, C5, and H3 originally targeting the Wuhan RBD, to bind more effectively to the Omicron RBD. A structure-based *in silico* affinity maturation pipeline was developed to enhance their binding affinities. The pipeline consists of three key steps: high-throughput *in silico* mutagenesis of complementarity-determining regions (CDRs), protein-protein docking for screening, and molecular dynamics (MD) simulations for assessment of the complex stability. A total of 741, 551, and 684 mutations were introduced in H11-H4, C5, and H3 nanobodies, respectively. Protein-protein docking and MD simulations shortlisted high-affinity mutants for H11-H4(6), C5(5), and H3(6). Further, recombinant production of H11-H4 mutants and Omicron RBD enabled experimental validation through Isothermal Titration Calorimetry (ITC). The H11-H4 mutants R27E, S57D, S107K, D108W, and A110I exhibited improved binding affinities with dissociation constant (K_D_) values ranging from ∼8.8 to ∼27 µM, compared to the H11-H4 nanobody K_D_ of ∼32 µM, representing a three-fold enhancement. This study demonstrates the potential of the developed *in silico* affinity maturation pipeline as a rapid, cost-effective method for repurposing nanobodies, aiding the development of robust prophylactic strategies against evolving SARS-CoV-2 variants and other pathogens.

**Graphical Abstract:** 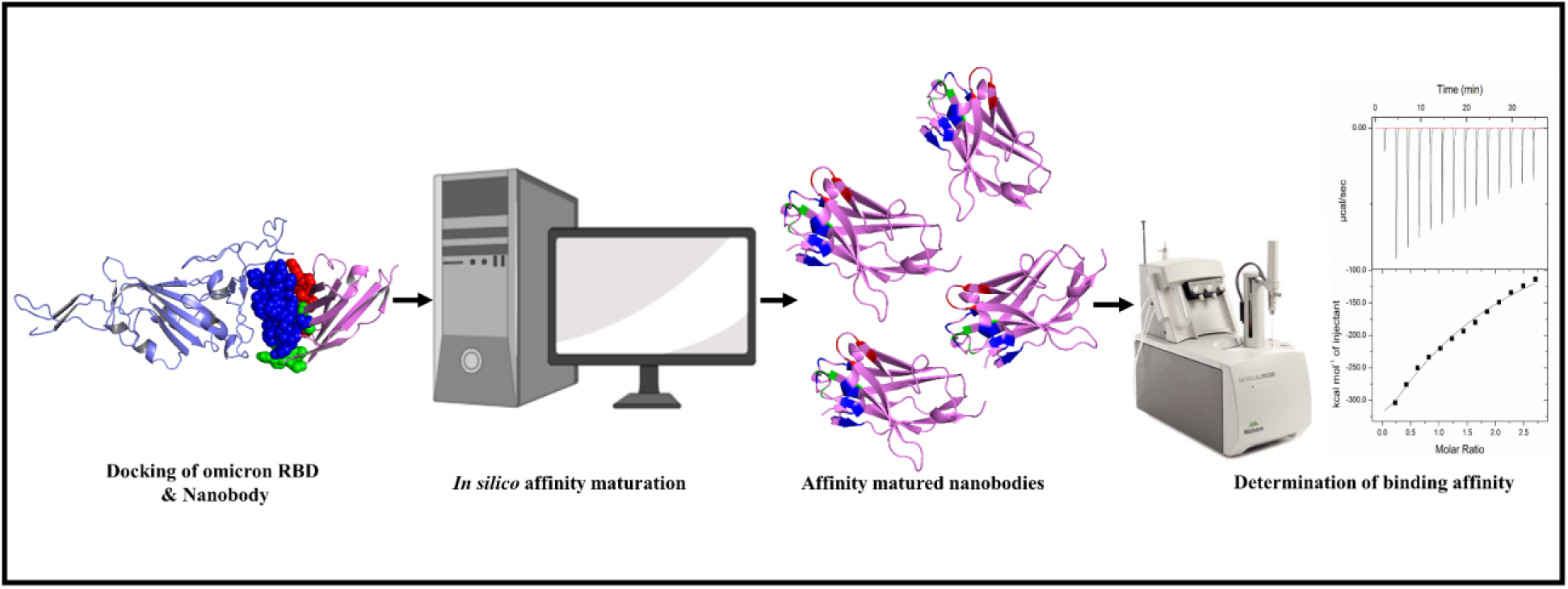

## 1. Introduction

The serendipitous discovery of heavy-chain only antibodies (HcAbs) in camelids has ignited the development of “third generation” antibodies, significantly expanding treatment options with the opening of new research avenues. Unlike conventional antibodies, these HcAbs lack light chains and consist solely of homodimeric heavy chains, circulating naturally in Camelidae and some sharks (1,2). The antigen-binding domain of these antibodies known as “variable heavy domains of heavy chains” (VHH) or “Nanobody” is a suitable candidate for straightforward engineering (3). This unique structure offers distinct advantages, including increased stability and solubility.

Nanobodies, with their compact size of ∼15 kDa, are more straightforward to clone, express, and purify efficiently, achieving high yields across bacterial, yeast, and mammalian systems. (2,4,5). Structurally, these are composed of a highly conserved framework region (FR), surrounded by three hypervariable loops known as complementarity-determining regions (CDRs) or paratopes. These CDRs are responsible for antigen recognition and binding specificity (1,6,7).

Nanobodies offer several advantages over conventional antibodies, including high specificity, thermodynamic stability, diverse paratopes, superior target engagement with the ability to access hidden or intracellular epitopes, and better tissue penetration (2). Underscoring the potency and efficacy of nanobodies, recently a therapeutic nanobody Caplacizumab has been approved as a treatment for an immune-mediated deficiency called thrombotic thrombocytopenic purpura (TTP) (8). Traditionally, nanobodies have been produced through the immunization of camelids.

Additionally, these can be successfully screened from large libraries (immune, naïve, and synthetic) through display technologies (phage, yeast, or ribosome) (5,9). However, a plethora of nanobodies generated by these methods do not achieve optimal affinity or specificity for their target antigens and require further *in vitro* affinity maturation (10–13). Methods such as mutational hotspot randomization, error-prone polymerase chain reaction (PCR), random affinity mutagenesis, and CDR walking were commonly used to enhance the affinity of antibodies (13–16). Several *in vitro* affinity maturation techniques utilizing directed evolution, diversify antibody sequences to select the best binders. These techniques effectively mimic the somatic hypermutation process observed in mammalian B cells for enhancing the affinity and specificity of the naturally occurring antibodies (10,17). Experimental screening of vast antibody libraries is not only costly and time-consuming but is also hindered by challenges such as low expression, poor solubility, and polyspecificity of screened antibodies (18,19). These difficulties emphasize the necessity for advanced alternative techniques in antibody development.

Computational design has emerged as a promising method for discovering and optimizing antibodies. It provides a cheaper and faster alternative to traditional *in vivo* and *in vitro* techniques, allowing better control over key biophysical properties such as stability and solubility (12,18). A complementary approach utilizing computational methods such as *ab initio* modeling, *in silico* random or site-directed mutagenesis, alanine scanning, energy calculations, and machine learning offers a new strategy for the generation of high affinity and stable binders, are more efficient than traditional methods (12,18,20,21). Structure-based antibody design leverages high-resolution structural data to guide mutations that enhance antibody-antigen interactions. However, their success depends on the presence of a high-quality structure of antibody-antigen and the availability of an algorithm capable of predicting the energetic impacts of mutations (18,21). The primary advantage of this approach lies in its precision; by focusing on the three-dimensional structure of the antibody-antigen interface, to predict how specific mutations will affect binding affinity and specificity. This allows for targeted modifications, rather than relying on random mutations, leading to more efficient optimization of antibody performance. Furthermore, the use of *in silico* tools, such as protein-protein docking, molecular dynamics (MD), and neural network-based algorithms, provides a powerful means to predict the energetic and stability changes caused by mutations. These predictive models streamline the process of affinity maturation, enabling the development of highly specific antibody fragments like fragment antigen-binding regions (Fabs), single-chain variable fragments (scFvs), and VHHs that can target a broad spectrum of antigens (12,18,20–23).

The first critical step in the life cycle of SARS-CoV-2 is the interaction of Spike-protein (S-protein) with the angiotensin-converting enzyme 2 (ACE2) receptor of the host cell, subsequently leading to membrane fusion and cell entry (24,25). Antibody-based therapies targeting the S-protein aim to neutralize the virus by disrupting its interaction with ACE2 or by preventing conformational changes in the S-protein (9,26,27). Similarly, nanobodies neutralize the virus either by directly competing with ACE2 for the same epitope (Ty1, H11-H4, C5, and H3) or by interacting with the non-receptor binding domain (RBD) of S- protein (F2, C1, and VHH72) and preventing the confirmational changes (28–31). Moreover, nanobodies are effective therapeutic or prophylactic agents that could be delivered through the intranasal route against SARS-CoV-2 (32,33). Notably, PiN-21, an aerosol-based nanobody for the prevention and treatment of SARS-CoV-2, provides evidence for a significant reduction in viral loads, prevention of lung damage, and virus induced pneumonia (34). RNA viruses, including SARS-CoV-2, frequently acquire mutations in their genomes (35). While lethal mutations in replication proteins are naturally filtered out by selection pressures, non-lethal mutations in structural proteins that improve viral fitness and survival tend to be preserved (36). The SARS-CoV-2 variant of concern (VOC), Omicron (B.1.1.529), carries 30 mutations in the S-protein, including 15 mutations exclusively in the RBD (37,38). Mutations in the S-protein affect the transmissibility of the virus and the efficacy of neutralizing antibodies generated through vaccination or previous infections (39). This dynamic interplay between mutation and immune response challenges the development of effective vaccines and therapeutics. As a key receptor-engaging protein, the RBD of SARS-CoV-2 is a promising target for suppressing and inhibiting virus infection. The COVID-19 pandemic sparked a surge in research focused on high-affinity nanobodies. Several studies on SARS-CoV-2 therapeutics have demonstrated the effectiveness of nanobodies that target the RBD of the S-protein(9,26,28–30,32–34,40–46).

In line with these studies, the present study reports a streamlined computational pipeline to generate high affinity and stable mutants of three nanobodies H11-H4, H3, and C5 that are reported to neutralize the Wuhan isolate of SARS-Cov-2 by directly targeting ACE2 binding site of RBD of S-protein. Focusing on the Omicron variant of SARS-CoV-2, an *in silico* nanobody engineering workflow is outlined here, which focuses on the generation of a library of high affinity and stable nanobodies through substitution mutations in the CDRs of these nanobodies. The crystal structure of the three nanobodies in complex with RBD of SARS-CoV-2 (PDB ID: 6ZBP, 7OAN, and 7OAP) served as a reference point for high-throughput *in silico* substitution mutagenesis. An *in house* library of mutants was generated and screened against Omicron RBD using protein-protein docking. MD simulation studies further refined the selection based on dynamic stability and interactions. To validate the *in silico* predictions, recombinant Omicron RBD and H11-H4 nanobody/mutants were produced and their binding affinities were directly measured using Isothermal Titration Calorimetry (ITC). Taken together, the present study provides an *in silico* pipeline for computational affinity maturation of established and verified nanobodies, paving the way for their development as therapeutics, diagnostics, and drug delivery systems.

## 2. Methodology

### 2.1 Software and hardware used in the study

Protein structures were obtained from the RCSB Protein Data Bank (PDB) (47), with their IDs mentioned as needed throughout the text. Protein-protein docking was performed using the High Ambiguity Driven protein-protein Docking (HADDOCK 2.4) webserver (48). Structural visualization, comparisons, and interaction studies were performed using PyMOL (49) and LigPlot+ (50). *In silico* affinity maturation was executed using mCSM-AB2 (51), DUET (52) and Deep DDG (53) web servers. Mutations in the nanobodies were introduced using the DynaMut (54) web server. MD simulations were carried out with the Gromacs 2022.2 suite (55) on an Ubuntu-based Linux workstation.

### 2.2 Computational affinity maturation and generation of the virtual library

Nanobodies can be categorized into two groups based on their interaction with the RBD. For instance, Cluster 1 nanobodies bind sideways to the RBD whereas Cluster 2 nanobodies target and cover the ACE2 binding site of RBD (56). Three nanobodies from cluster 2, H11-H4, C5, and H3 that were previously reported to interact with the RBD of Wuhan isolate of SARS-Cov-2 were selected for the present study (29,30). The 3D crystal structures of Omicron RBD and nanobodies were obtained from RCSB-PDB using PDB ID: 7WBP, 6ZBP, 7OAN, and 7OAP (29,30,57). The structures were prepared by removal of water and other co-crystallized ligands/proteins using PyMOL.

These nanobodies were then docked with the Omicron RBD using HADDOCK 2.4. The interacting residues (active residue) reported to be located at the interface of the RBD of Wuhan isolate SARS-Cov-2 and nanobody complex (PDB ID: 6ZBP, 7OAN, and 7OAP) were used as constraints to generate AIRs (Ambiguous Interaction Restraints) during protein-protein docking, keeping other parameters as default. The top-ranked Omicron RBD and nanobody complexes, based on HADDOCK 2.4 score and cluster size, were selected and analyzed to study the detailed interaction patterns, using the DIMPLOT module of LigPlot+ v2.2.4 for two-dimensional (2D) representation and were visualized using PyMOL for 3D visualization.

The Omicron RBD and nanobody complexes generated through the HADDOCK 2.4 webserver were fed into the mCSM-AB2, DUET, and the Deep DDG webserver for *in silico* mutagenesis to predict changes in the binding affinities and stability upon mutations in CDR loops of nanobodies. Through the *in silico* mutation module of mCSM-AB2, each residue present in the CDR loops of nanobodies was mutated into 19 amino acids other than itself, and the resulting changes in binding affinities were calculated as ΔΔG (Gibbs free energy; kcal/mol), with ddG > 0 indicating increased affinity and ddG < 0 indicating decreased affinity compared to wild type (51). The selection of stabilizing mutations in nanobodies was done using an integrated approach of DUET and the Deep DDG webserver. Deep DDG, an online neural network-based method, is used to calculate the effects of point mutations in a protein by predicting the changes in its stability, with ddG > 0 indicating increased stability and ddG < 0 indicating decreased stability compared to wild type. (52,53). Utilizing this approach, an *in house* library comprising high binding and stable mutations in nanobodies H11-H4, C5, and H3 was generated.

### 2.3 Interaction assessment and selection of best mutants through docking studies

The mutations identified through computational affinity maturation were introduced in nanobodies H11-H4, C5, and H3 using DynaMut web server. It is a user-friendly web server that predicts how mutations alter a protein’s conformation and movement by analyzing vibrational entropy changes (54). The resulting mutated structures were then docked against the Omicron RBD using HADDOCK 2.4, employing the same constraints as used in the previous docking. These docked complexes were screened based on docking scores and favorable interactions compared to the native nanobody and RBD complex. A detailed comparative analysis of ionic, polar, and non-polar interactions was done using DIMPLOT and PyMOL.

### 2.4 Analyzing nanobody and RBD complexes via MD simulations

To study and validate the stability of the mutant nanobody: RBD complexes at the atomic level, MD simulation studies were performed for the complexes selected after docking studies. The simulation was executed using the GROMOS96 43a1 force field in the GROMACS 2019.5 suite. The systems were solvated using a simple point charge (SPC) water model in a cubic box with 1 nm marginal radii. Counter ions Na^+^ and Cl^-^ were added to maintain the overall neutrality of the system by replacing the water molecules. To minimize steric clashes and steepest descent, an energy minimization algorithm was used for 50,000 iterations steps with the cut-off of 10.0 kJmol-1. Particle Mesh Ewald (PME) with 1.6 Å Fourier grid spacing was used for the calculation of Long-range electrostatics and Coulomb interactions were calculated within a cut-off radius of 12 Å (58). The short-range forces were calculated with a minimum cutoff set to 12 Å using a verlet cut-off scheme. Followed by a two-step equilibration phase of, a constant number of particles, volume, and temperature (NVT) for 1 ns and a constant number of particles, pressure, and temperature (NPT) for 1 ns at a temperature of 300 K, using V-rescale temperature coupling method and Berendsen coupling method, respectively (59). Finally, the leap-frog algorithm was employed to carry out a 100 ns MD run for the system with an integration time frame of 2 fs, and the trajectories were generated after every 10 ps (60). The relative RMSD (Root Mean Square Deviation) values with respect to the initial reference trajectory for RBD and nanobody complexes were determined using g_rms. Analysis of the average number of intermolecular H-bonds at the interface of RBD and nanobody complexes was done using g_hbond within the GROMACS suite for the 100 ns simulation. The trajectories of MD run were analyzed using g_dssp, to determine the secondary structure changes in nanobodies.

### 2.5 Expression, purification and site-directed mutagenesis in nanobody

The H11-H4 nanobody gene, previously cloned into the pET28c expression vector (61), served as the template for SDM. Single amino acid substitutions within the CDRs of the nanobody were introduced using overlapping primers, yielding a library of six distinct nanobody variants **(Supplementary Table 1).** Following amplification with enzyme *Phusion High-Fidelity DNA polymerase* (Thermo Scientific), the parental plasmid was digested with *DpnI*, exploiting its methylation sensitivity. The digested product was transformed into competent *E. coli* XL-1 Blue cells and plated on Luria Bertani (LB) agar (HiMedia, India) with kanamycin and chloramphenicol for selection. Plasmid from selected colonies was then isolated and the mutations were confirmed by Sanger sequencing.

Expression and purification of H11-H4 (native) and its mutants were done following the same protocol as described earlier (61). Briefly, the recombinant plasmid was transformed into chemically competent *E. coli* Rosetta (DE3) cells for protein expression, with selection on LB agar plates containing kanamycin (50 μg/mL) and chloramphenicol (35 μg/mL). A single colony was used to inoculate a 10 mL LB broth primary culture with the same antibiotics and grown overnight in a shaking incubator at 37 °C. This primary culture was then used to inoculate a 1 L LB broth for protein production, grown at 37 °C until the mid-log phase (OD₆₀₀ ≈ 0.4). Protein expression was induced with 0.2 mM isopropyl-β-d-1-thiogalactopyranoside (IPTG), and the culture was incubated at 16 °C for 20 h. Later, cells were harvested by centrifugation at 4000 × g for 10 mins and the pellet was resuspended in lysis buffer for disruption using a French press (Constant Systems Ltd, Daventry, England). After centrifugation, the clarified supernatant was loaded onto a pre-equilibrated Ni-NTA affinity chromatography column. Unbound components were washed out with increasing concentrations of imidazole and eluted fractions containing the purified nanobody were pooled and dialyzed overnight at 4 °C. Purity of nanobodies (∼15 kDa) were confirmed using Sodium Dodecyl Sulphate-Polyacrylamide Gel Electrophoresis (SDS-PAGE).

### 2.6 Cloning, expression, and purification of Omicron RBD

The gene sequence encoding the Omicron (B.1.1.529) RBD was retrieved from GISAID EPI_ISL_6640916 and the codon-optimized sequence was synthesized by GeneScript (Biotech Desk Pvt. Ltd; India). Expression and purification of the Omicron RBD followed the previously described protocol (61). Briefly, the RBD gene was amplified using forward primer 5’-TACGTACCAACTGAATCTATCGTTAGATTCCC-3’ and reverse primer 5’-CCTAGGCTAATGATGATGATGATGATGATT-3’ and then cloned into the pPIC9K vector between *AvrII* and *SnaBI* restriction sites. The clone confirmed by Sanger sequencing was linearized with *SalI* and transfected into competent *Pichia pastoris* cells via electroporation. After transfection cells were grown for 2-3 days on histidine deficient media for the selection of successfully transfected colonies. Positive colonies were screened with an increasing gradient of gentamycin to enrich high protein expressing colonies. Subsequently, selected colonies were evaluated for small-scale protein expression, and the highest-yielding colony was chosen for large-scale protein production. Later, the selected colony was cultured in Buffered Glycerol-based Medium (BMGY) at 28 °C with shaking until an OD₆₀₀ of ∼ 2-6 was reached. The cells were harvested and resuspended into a Buffered Methanol-complex Medium (BMMY) with 1% methanol for RBD protein expression. The culture was maintained at 28 °C and induced with 1% methanol for 4 days. Following the induction period, the culture supernatants containing RBD protein were collected and purified using Ni-NTA affinity chromatography. The eluted fractions were pooled and dialyzed overnight at 4 °C. The purity of RBD (∼45 kDa) was confirmed using SDS-PAGE.

### 2.7 ITC-based analysis of nanobody mutant and Omicron RBD interaction

The binding affinities of the nanobody and its mutants towards the Omicron RBD were determined using ITC (Malvern, Northampton, 248 MA), as per the protocol detailed previously (61). In brief, the experiment consisted of 14 injections at a stirring rate of 750 rpm and a temperature of 25 °C. Both proteins were diluted in 1 X PBS buffer of pH 7.4, with the RBD concentration set to 2.5 μM, and the nanobody concentration in the syringe was kept at 50 μM. Prior to the experiments, all samples were gently vacuum-degassed. The experimental setup consisted of an initial injection of 0.5 μL of nanobody into a reaction cell containing 350 μL of RBD, followed by 13 additional injections of 2.9 μL each, with 150 sec intervals between each injection. Following subtraction control (buffer), ITC data were fitted to a single-site binding model to calculate the dissociation constant (K_D_), enthalpy (ΔH), and entropy (ΔS) for the binding using MicroCal Analysis Software and Origin 7.0.

## 3. Results

### 3.1 Mutations in the S-protein altered the nanobody binding pattern

The Omicron variant of SARS-CoV-2 manifests a total of 30 substitutions, 6 deletions, and 3 insertions exclusively in the S-protein. A hot-spot site for mutations within the S-protein is the RBD, which is reported to have 15 substitution mutations (39,62). These mutations have enabled the Omicron variant to evade immune responses from prior infections or vaccinations, rendering established neutralizing antibodies ineffective and leading to numerous breakthrough cases in vaccinated individuals (39,63–66). Unlike, other prototypic variants with a hidden RBD, the Omicron variant has a stable open S-protein with a more exposed RBD (39,67). Therefore, neutralizing antibodies and nanobodies targeting the Omicron RBD could help in reducing the viral spread. Here, three cluster 2 nanobodies H11-H4, C5, and H3 were selected and docked with the Omicron RBD using HADDOCK 2.4. The best docked cluster was selected based on cluster size, docking score, and interaction patterns **(Table 1**). These nanobodies bind to the Omicron RBD similarly to the early pandemic viruses (Wuhan isolate). However, comparing the interactions in the RBD of Wuhan isolate SARS-Cov-2 and nanobody complexes (PDB IDs: 6ZBP, 7OAN, 7OAP) to those observed in this study revealed significant differences. To access the effect of RBD mutations on the RBD: nanobody interaction in detail the interaction present in Omicron RBD and nanobodies (H11-H4, C5, and H3) docked complexes were compared to the complexes of RBD (Wuhan isolate SARS-Cov-2) and nanobodies (PDB IDs: 6ZBP, 7OAN, and 7OAP) **(Supplementary Figure 1)**. Analysis of docked complexes in the DIMPLOT and PyMOL reveled the disruption of many polar and hydrophobic contacts and the formation of new bonds in the Omicron RBD and nanobody complexes **(Figure 1 and Supplementary Figure 1)**.

**Figure 1:**
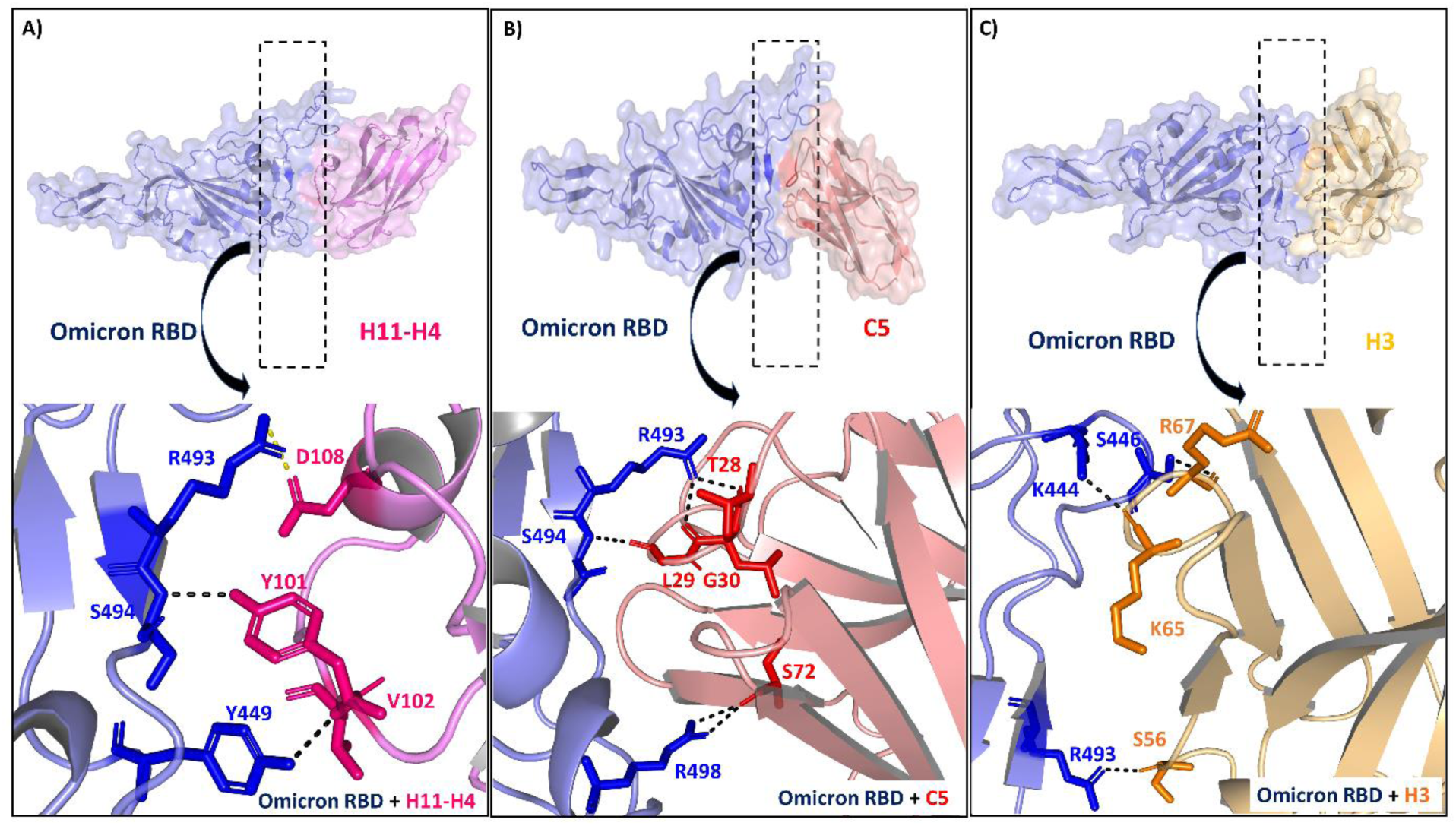
Structural representations of the interactions between the Omicron RBD and nanobodies. A) Surface representation of Omicron RBD (slate) and H11-H4 (violet) nanobody complex with the zoomed in view of H-bonded interaction present at the interface of RBD and H11-H4 nanobody. B) Surface representation of RBD (slate) and C5 (red) nanobody complex with the zoomed in view of H-bonded interaction present at the interface of RBD and C5 nanobody. C) Surface representation of RBD (slate) and H3 (orange) nanobody complex with the zoomed in view of H-bonded interaction present at the interface of RBD and H3 nanobody key interacting residues are highlighted and labeled, with H-bonds (black) and Salt bridges (yellow) depicted as dashed lines. The complexes highlight critical contact points contributing to the binding affinity and stability of the nanobody-RBD complexes.

**Table 1:**
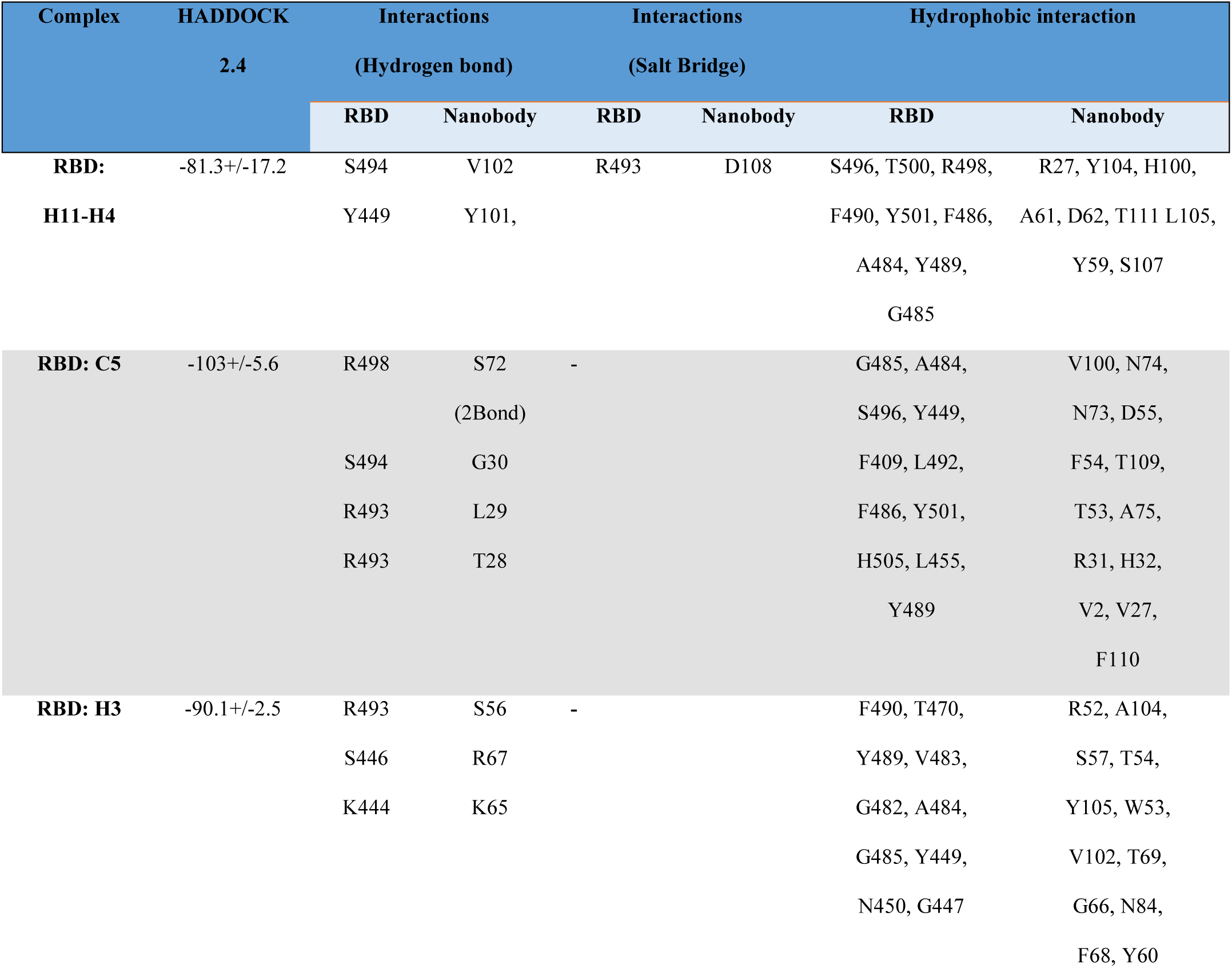
Comparative analysis of interactions present in docked complexes of Omicron RBD: nanobodies (H11-H4, C5, and H3) generated through HADDOCK 2.4. List of residues involved in electrostatic polar and nonpolar interactions identified using Dimplot.

The Q493R mutation in Omicron RBD stabilizes its docked complex with H11-H4 nanobody by forming a new salt bridge with D108 of the H11-H4, however, its H-bond with Y101 and Y104 of H11-H4 nanobody was lost. Moreover, the mutation E484A in the Omicron RBD led to the disruption of critical bivalent salt linkage with R52 and one H-bond with S57 of H11-H4 nanobody. A new hydrophobic interaction with Y104 and L105 of H11-H4 and F490 of Omicron RBD was observed in the complex of Omicron RBD: H11-H4 complex, though its H-bond with Y104 of H11-H4 nanobody was lost. The ACE2-interacting residue of RBD, Y449 formed a new H-bond with V102 of H11-H4 nanobody. A loss of one H-bond for S494 was observed in the complex of Omicron RBD: H11-H4 **(Figure 1A and Supplementary Figure 1)**. The Omicron RBD mutations E484A and N501Y, resulted in the loss of H-bonds with R31 and N73 of C5 nanobody respectively **(Figure 1B and Supplementary Figure 1)**. However, after mutation Q498R two new H-bonds were observed between R498 and S72 of the C5 nanobody. Additionally, after mutation Q493R, the mutated residue R493 forms two new H-bonds with T28 and L29 of C5 nanobody **(Figure 1B and Supplementary Figure 1)**. The complex of H3 nanobody with RBD (Wuhan isolate SARS-Cov-2) formed a total of ten H-bonds at the interface (PDB: 7OAP), however, only three H-bonds were observed in the docked complex of Omicron RBD: H3 **(Figure 1C and Supplementary Figure 1**). Owing to the E484A mutation in Omicron RBD, the loss of four H-bonds was observed in the Omicron RBD: H3 complex. Mutation Q498R resulted in the disruption of its H-bond with A104, instead, the formation of a new H-bond with S56 of H3 nanobody was observed. K444 and the mutated residue S446 (G446S) form two new H-bonds with residue K65 and R67 of H3 nanobody respectively **(Figure 1C and Supplementary Figure 1)**. Details of electrostatic, polar, and nonpolar interactions present in the docked complexes of Omicron RBD and nanobodies (H11-H4, H3, and C5) were explained in **(Table 1).**

### 3.2 Generation of *in house* virtual library and *in silico* affinity maturation

The Omicron RBD and nanobody complexes for H11-H4, H3, and C5 nanobodies generated using HADDOCK 2.4 served as input for the *in silico* affinity maturation software mCSM-AB2, DUET, and Deep DDG to predict high-affinity and stable mutations in nanobodies. The CDR loops of the nanobodies were targeted for multiple substitution mutations in residues of H11-H4 (CDR1: G26-M34, CDR2: A50-Y59, CDR3: A97-Y116), C5 (CDR1: G26-A33, CDR2: R52-T58, CDR3: A97-F110) and, H3 (CDR1: G26-S33, CDR2: M51-T58, CDR3: A97-S116). This resulted in a virtual library comprising mutations for H11-H4 (741), C5 (551), and H3 (684). The mCSM-AB2 software predicted the effect of mutations in nanobodies, as a function of binding affinity, and an integrated approach of DUET and Deep DDG software was applied, to assess the effect of mutations on the stability of Omicron RBD and nanobody complex. Further common mutations were filtered to narrow down the total number of mutations by selecting those mutations that have the predicted ΔΔG > 0 indicating increased affinity and stability **(Figure 2)**.

**Figure 2:**
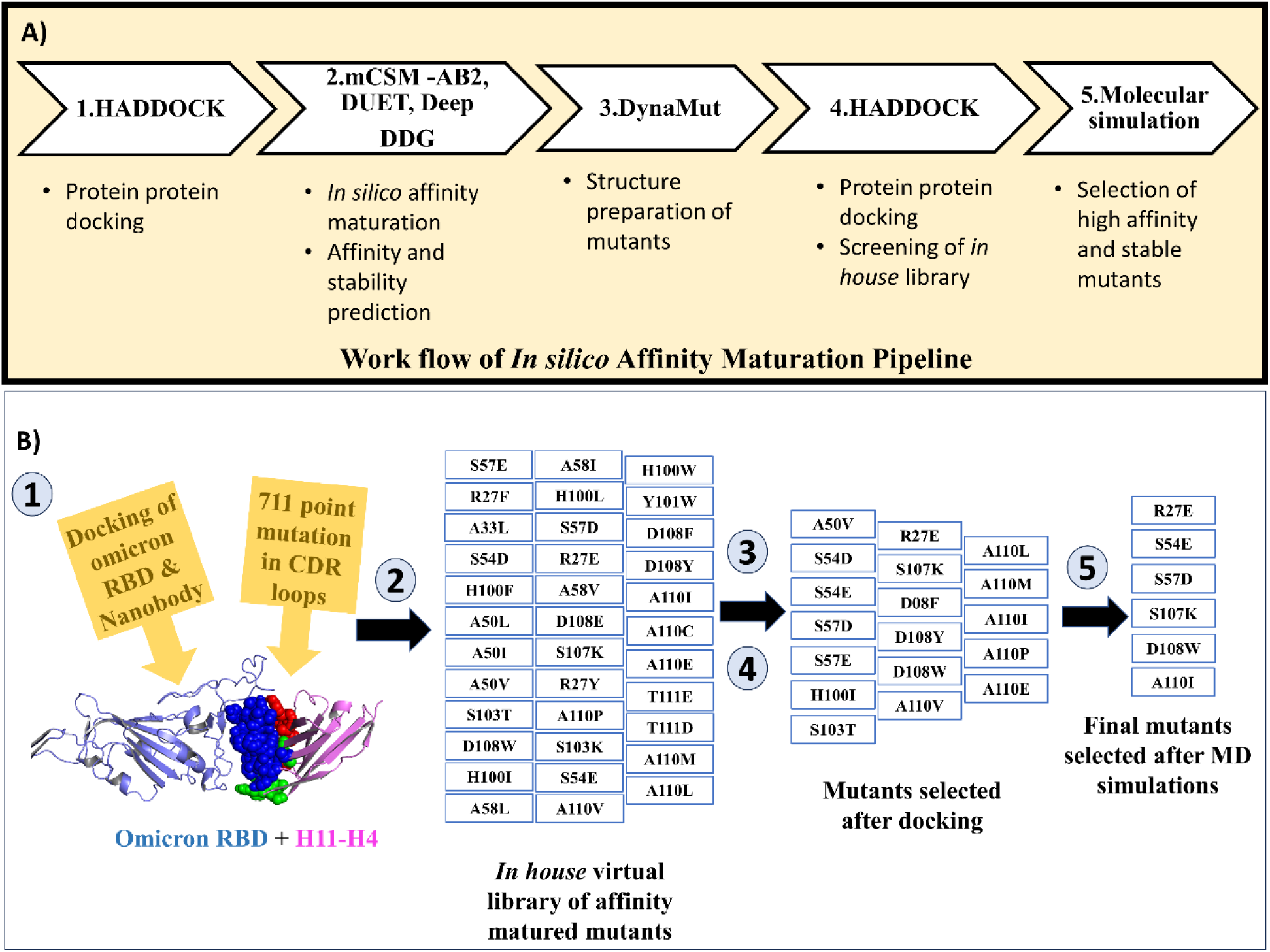
*In Silico* Affinity Maturation Pipeline for Optimizing Nanobody Binding to Omicron RBD. A) Workflow of the *in silico* affinity maturation pipeline used to enhance nanobody binding affinity to the Omicron RBD. The process begins with protein-protein docking using HADDOCK (1), followed by *in silico* affinity maturation and stability prediction using mCSM-AB2, DUET, and DeepDDG web serves (2). Mutants are prepared structurally using the DynaMut web server (3), and then protein-protein docking is repeated for screening the *in house* library of mutants (4). Finally, high-affinity and stable mutants are selected through molecular simulations (5). B) Detailed steps in the workflow for H11-H4 nanobody: (1) Initial docking of Omicron RBD with nanobody H11-H4. (2) Generation of a virtual library with 711-point mutations in the CDR loops. (3) Screening of mutants post-docking to identify those with improved binding. (4) Selection of mutants after docking. (5) Final selection of high affinity and stable mutants post-MD simulations. The Omicron RBD is shown as cartoon (slate), while the nanobody H11-H4 is shown in purple with residue of CDR1 (green), CDR2 (red), and CDR3 (blue) shown as Spheres.

The predicted ΔΔG values were used to decode the criteria for predicting high-affinity and stable mutations. Interestingly, mCSM-AB2 software preferred aromatic residues Tryptophan (W), Tyrosine (Y), Phenylalanine (F) and charged residues Glutamate (E), Aspartate (D), Arginine (R), Lysine (K), Histidine (H) for substitutions (68). The highest ΔΔG values were predominantly for aromatic residues, with A110Y in H11-H4, G99W in C5, and S33W in H3 nanobody **(Figure 3)**. In contrast, the integrated approach using DUET and Deep DDG software showed a preference for non-polar residues Leucine (L), Valine (V), and Isoleucine (I) residues. The highest ΔΔG values were observed for Q98L in H11-H4, and E111L in H3, but D106C had the highest value in C5 **(Figure 3).**

**Figure 3:**
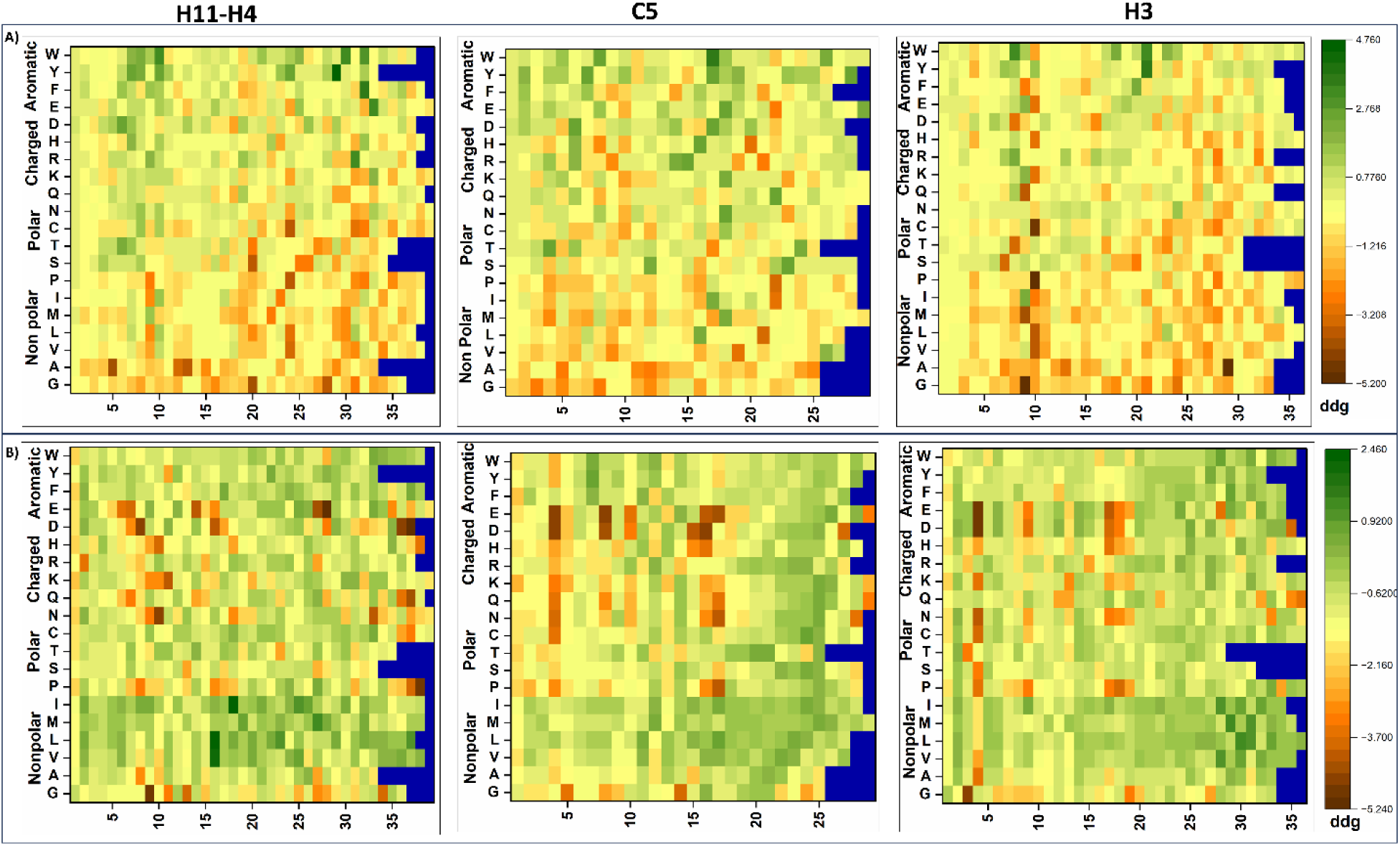
Heat maps illustrating the ΔΔG values for residue substitutions predicted by mCSM-AB2 (panel A) and Deep DDG (panel B). In each heatmap row represents a specific substitution (Y-axis) across different positions (X-axis) in three different sequences: H11-H4 (left two heat maps), C5 (middle two heat maps), and H3 (right two heat maps). The color scale indicates the ΔΔG values, with green representing high affinity/stabilizing mutations and brown representing low affinity/destabilizing mutations. mCSM-AB2 shows a general preference for aromatic and charged substitutions, as evidenced by the clustering of stabilizing mutations (green) in these regions. Deep DDG shows a general preference for nonpolar and aromatic substitutions, as evidenced by the clustering of stabilizing mutations (green) in these regions. Differences between the two prediction tools highlight varying propensities for certain substitutions across positions.

After exhaustive single-point virtual mutagenesis within the CDRs of nanobodies, 35 mutations each for H11-H4 and H3 nanobodies, and 26 mutations for the C5 nanobody were identified as stable affinity enhancing mutations. These were selected for further detailed docking studies **(Supplementary Tables 2, 4, and 6)**.

The mCSM-AB2 and Deep DDG software identified affinity enhancing and stable mutations in the H11-H4, C5, and H3 nanobodies. Comparative analysis of mutational groups reveals distinctive residue compositions and substitution patterns within nanobodies. The H11-H4 nanobody was characterized by a composition rich in charged residues and aromatic residue, with 14 out of 35 substitutions converting A residues to other amino acids **(Figure 4 and Supplementary Table 2)**. In contrast, C5 exhibits a deficit in aromatic residues and predominantly substitutes nonpolar residues in 13 out of 26 substitutions, with limited involvement of aromatic substitutions **(Figure 4 and Supplementary Table 4)**. H3, distinguished by its high proportion of charged residues, shows moderate aromatic residue content and substitution patterns converting nonpolar residues. Substitution trends indicate that H11-H4 and H3 exhibit a notably higher proportion of charged residues **(Figure 4 and Supplementary Table 6)**. These findings provide significant insights into the preferences for mutational residue types, which may be influenced by the specific residue located at the interface between nanobody and Omicron RBD complexes.

**Figure 4:**
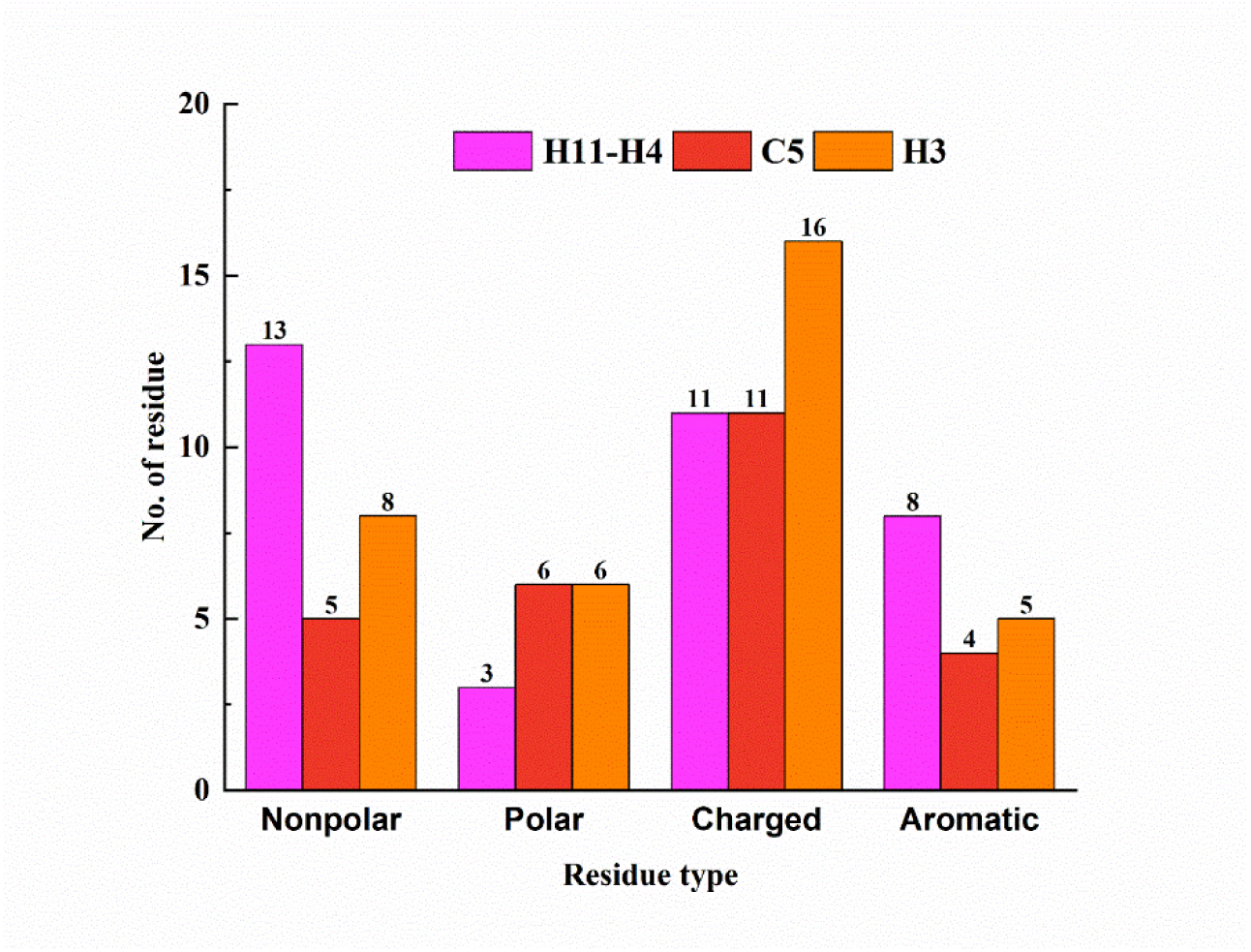
Distribution of residue types in the selected mutations of H11-H4, C5, and H3 using mCSM-AB2 and Deep DDG software. The bar graph shows the number of nonpolar, polar, charged, and aromatic residues for each group. Nanobody H11-H4 (pink) has 13 nonpolar, 3 polar, 11 charged, and 8 aromatic residues. Interestingly out of 35 substitutions, 14 substitutions were A to other residues, whereas only one substitution was from any aromatic residue to Tyr residue. Nanobody C5 (red) has 5 nonpolar, 6 polar, 11 charged, and 4 aromatic residues. For C5, out of 26 substitutions 13 substitutions were from any nonpolar to other residues, and 3 substitutions form any aromatic residue to other charged or aromatic residues. Nanobody H3 (orange) has 8 nonpolar, 6 polar, 16 charged, and 5 aromatic residues. Here, out of 35 substitutions, 9 substitutions were from nonpolar to other residues, whereas only three substitutions were from within aromatic residues. This visualization highlights the varying compositions of residue types among the three groups of mutations.

### 3.3 Selection of high affinity mutants using docking and MD simulation

High-affinity substitution mutations identified from the first round of screening were introduced in nanobodies using DynaMut software. The resulting 3D structures of the mutated nanobodies were subsequently docked with the Omicron RBD using HADDOCK 2.4. The docked clusters were analyzed, and the top-scoring cluster, exhibiting a higher number of favorable interactions, was selected. For further studies, 18 mutants of H11-H4, 14 mutants of H3, and 8 mutants of C5 were shortlisted. **(Supplementary Tables 2, 4, and 6)**. For selected RBD and nanobody complexes MD simulations were carried out to understand the structural changes at the atomic level after mutations. The MD runs trajectories were analyzed using XMGRACE. RMSD determines the conformational changes occurring in the protein backbone during the simulations, indicating the dynamic stability of the complex. The average RMSD for the RBD and native nanobody H11-H4, C5, and H3 complexes were 0.61 nm, 0.42 nm, and 0.5 nm respectively. For the RBD and mutant nanobody complexes, the average RMSD values were in the ranges from 0.36 nm to 0.71 nm for H11-H4, 0.42 nm to 0.75 nm for C5, and 0.35 nm to 0.77 nm for H3 **(Figures 5A and 5C and Supplementary Figures 2A and 2C)**. The H-bonds present at the interface of the protein-protein complex are critical factors affecting the stability and integrity of the complex. The average number of H-bonds present at the interface RBD and native nanobody H11-H4, C5, and H3 complexes were 9.5, 7.4, and 7.71 respectively. The average number of H-bonds for RBD and mutant nanobody complexes were in the range of H11-H4 (5.1 - 11.1), C5 (7.4 - 9.5), and H3 (3.2 - 10.3) **(Figures 5B and 5D and Supplementary Figures 2B and 2D)**. RBD mutant nanobodies complexes having average RMSD values and average H-bond numbers comparable to RBD and native nanobodies were selected for further studies **(Supplementary Tables 3, 5, and 7).**

**Figure 5:**
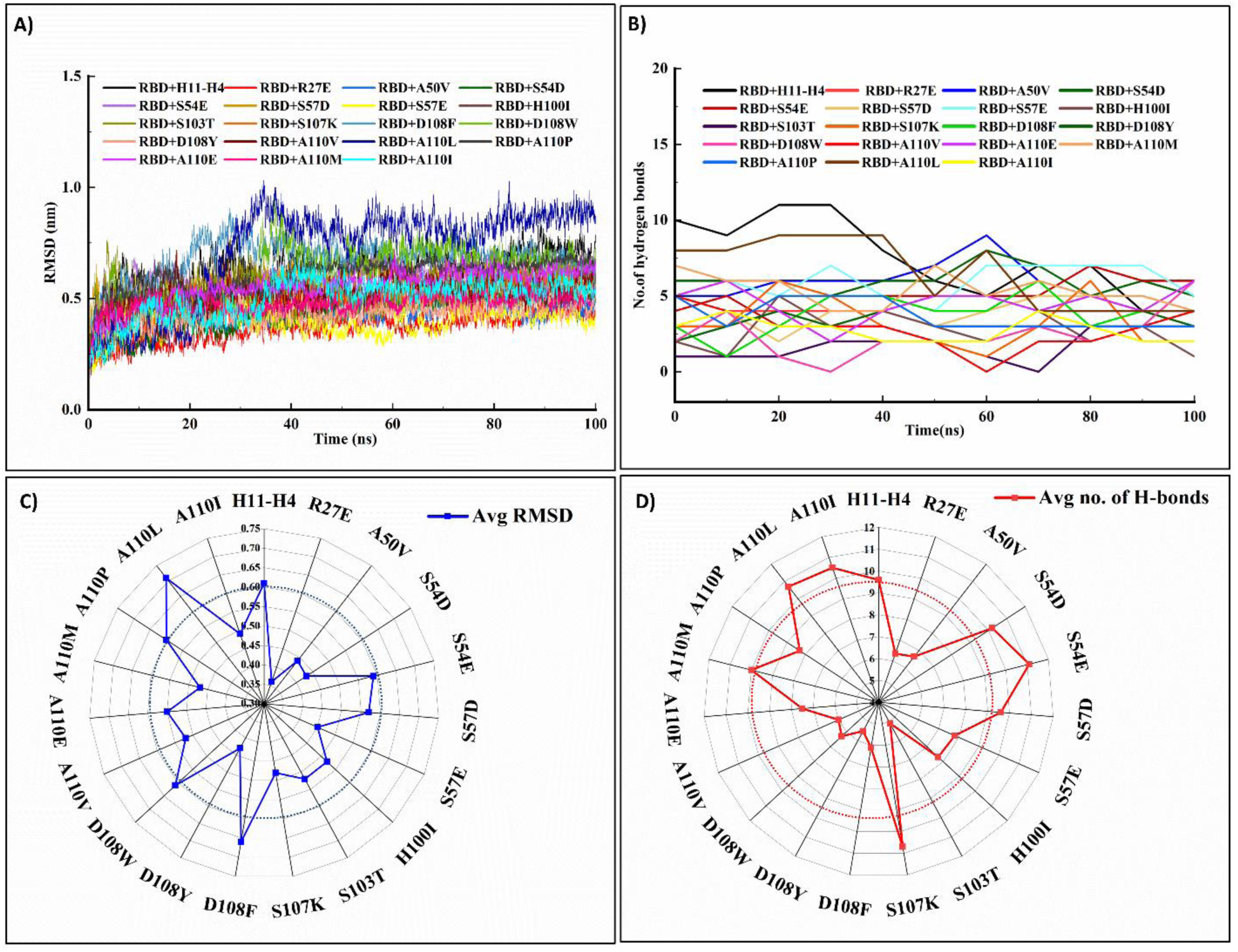
MD simulations studies determined the structural stability and interaction strength of Omicron RBD: mutant nanobody complex. A) RMSD profiles of RBD complexes with various nanobody mutants over 100 ns MD simulations. B) Number of hydrogen bonds formed between RBD and nanobody mutants over time. C) Radar plot depicting the average RMSD values for each nanobody mutant complex. D) Radar plot showing the average number of hydrogen bonds for each nanobody mutant complex.

### 3.4 Diverse mutational landscape affecting interactions of H11-H4 mutants and Omicron RBD

Following MD simulation, studies were focused primarily on the H11-H4 nanobody and its mutants. Among the 18 selected mutations of H11-H4, 7 involved substitutions to charged residues, 6 to nonpolar residues, 3 to aromatic residues, and 2 to polar residues. These mutations were distributed across the CDRs, with 1 in CDR1, 5 in CDR2, and 12 in CDR3 **(Figure 6A)**. The high number of mutations in CDR3 is attributed to its length and its role as the primary contact surface with the RBD. Detailed analysis of the CDRs of the H11-H4 nanobody elucidated its structural features and interaction capabilities, offering insights into amino acid composition differences and similarities that underlie the binding mechanisms of nanobody mutants with RBD. The residue of RBD in the 4 Å cutoff of RBD and H11-H4 complex were K478, A484, G485, F486, Y489, F490, Y449, F453, R493, S494, Y495, S496, R498, and Y501 **(Figure 6B)**. Aromatic residues constituted half of the interface, with the remainder composed of three charged residues, two polar residues, and two nonpolar residues **(Figure 6B)**. As per the shape and nature complementarity, antibody and antigen complex interface exhibited the highest propensity for aromatic residues such as tyrosine and tryptophan (69,70). Strikingly, during *in silico* affinity maturation of the H11-H4 nanobody, there is a discernible trend towards substituting nonpolar residues with charged and aromatic residues **(Figure 4)**, establishing the fact that the binding ability of nanobody mutants to RBD is significantly influenced by the propensity and specific positioning of certain amino acids within the CDR loops. Additionally, some mutations in the Omicron RBD were detrimental to the binding of H11-H4 and its mutants. Such as, the E484A mutation disrupts the bivalent salt bridge essential for complex stability (30,41,61). A similar residue distribution analysis was performed for the Omicron RBD and ACE2 complex (PBD: 7WBP). In this complex, RBD in the 4 Å cutoff of ACE2 were Y449, Y453, F486, N487, G476, A475, N477, Y489, F456, S494, R493, S496, R498, T500, F501, G502, and H505 **(Figure 6C).**

**Figure 6.**
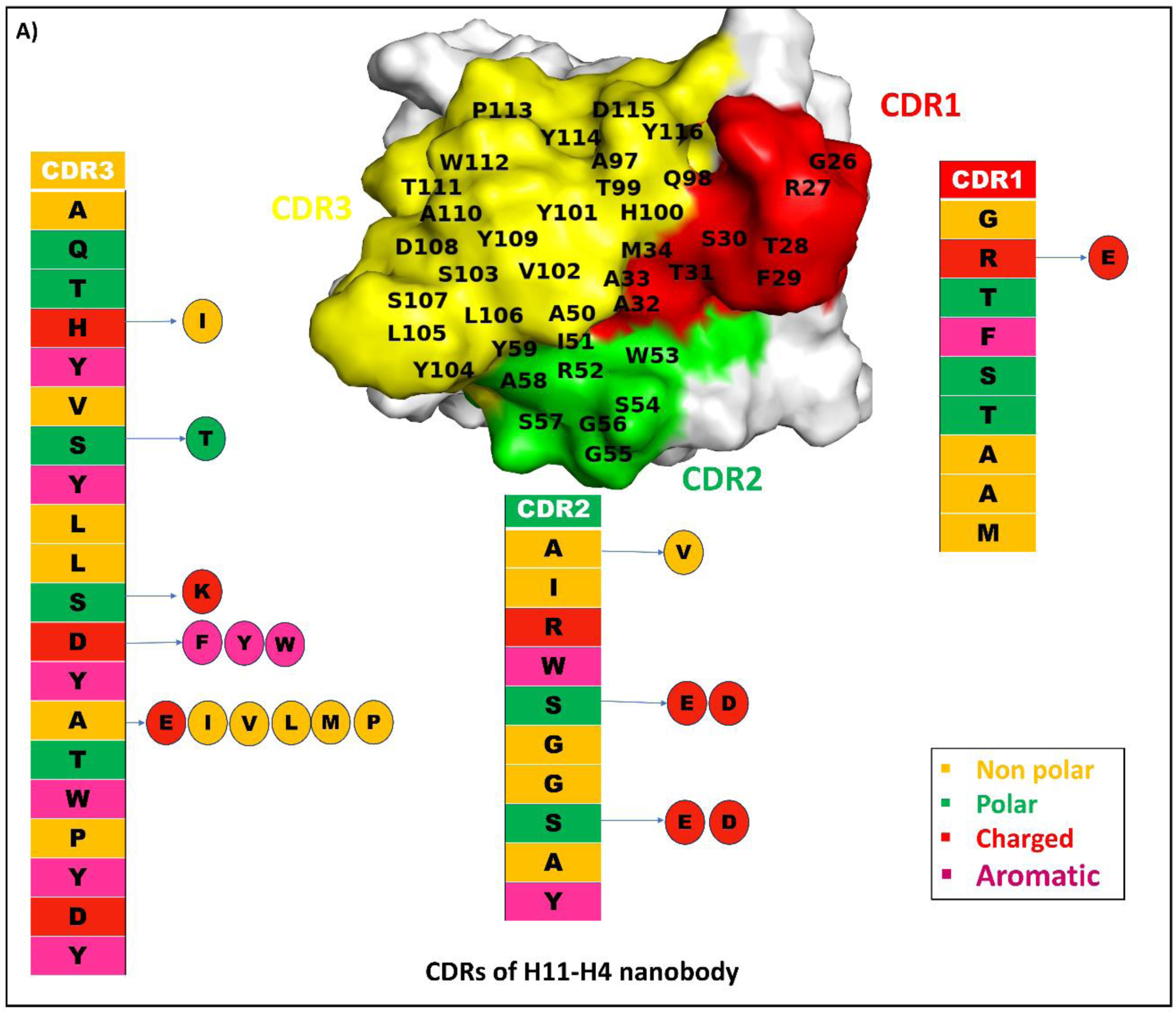
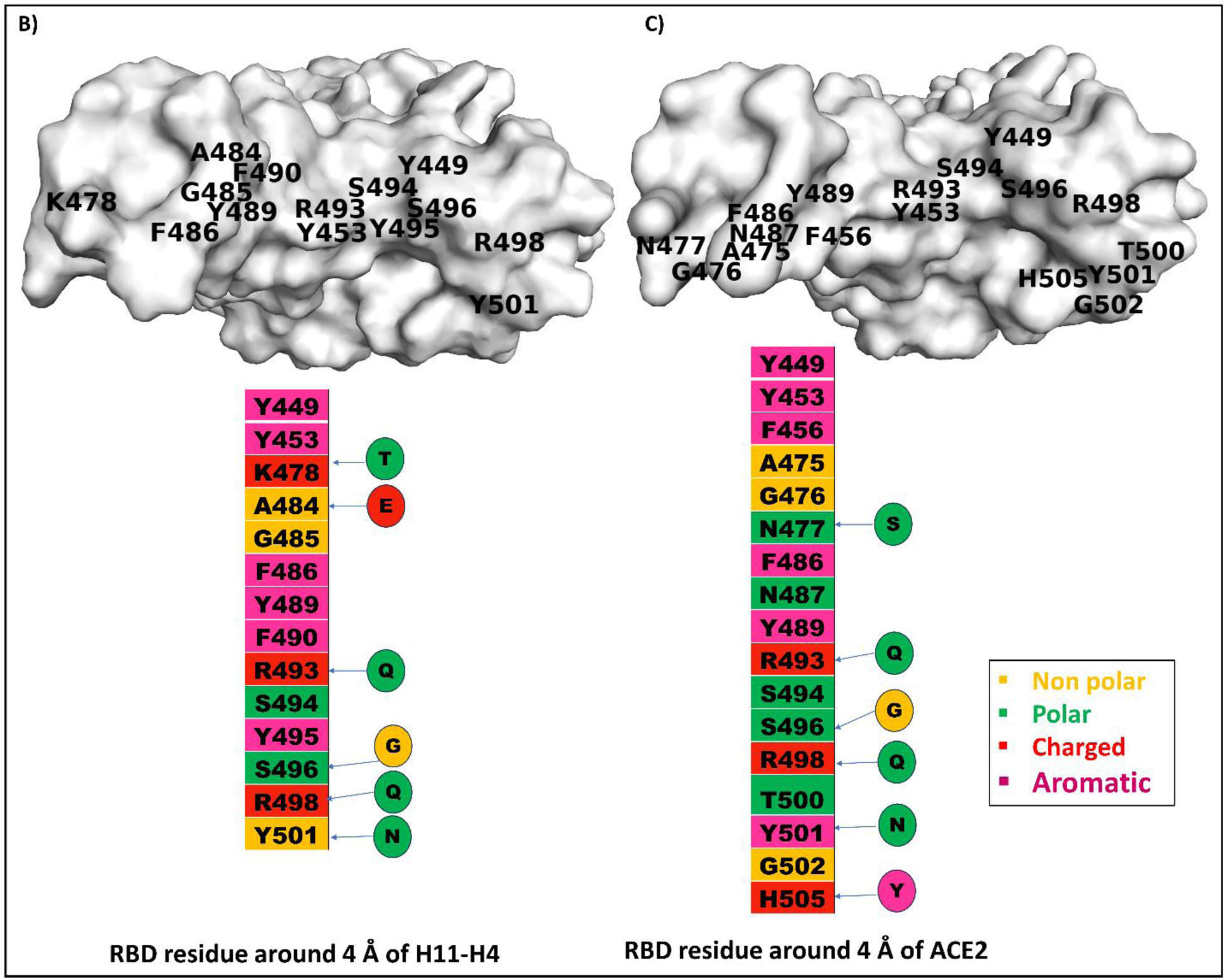
Amino acid distribution in the Paratope of H11-H4 nanobody and comparative analysis of residue distribution at the binding interface of ACE2 and H11-H4 on Omicron RBD. A) Surface view of H11-H4 nanobody showing the amino acid composition within the CDR1 (red), CDR2 (green), and CDR3 (yellow) of the H11-H4 nanobody. The vertical columns showcase the residue type of CDRs involved in the binding interactions. The arrow points out the mutations enclosed in circles that were selected in particular CDRs. B) Surface view of RBD showing the amino acid within the 4 Å of the ACE2 binding site of Omicron RBD (PBD: 7WBP). C) Surface view of RBD showing the amino acid within the 4 Å of the H11-H4 binding site of RBD. The vertical columns showcase the residue type of ACE2 and H11-H4 CDRs involved in the binding interactions. The arrow points out the mutations enclosed in circles in the Omicron variant.

Interestingly, over half of these residues were aromatic or charged, forming the key interaction interface with ACE2 **(Figure 6C).** It was reported that mutations in Omicron RBD particularly N501Y, Q493K/R, and T478K, augment the binding between the RBD and ACE2 by creating unique interaction patterns (62,64,71–74). Insights gained from the above analysis were instrumental in guiding the selection of H11-H4 nanobody mutants for further studies. One mutation from CDR1, two mutations from CDR2, and three mutations from CDR3 were chosen for further analysis, incorporating all CDRs and including polar, non-polar, charged, and aromatic residue substitution in the selection criteria.

### 3.5 Interaction studies elucidated binding characteristics of H11-H4 mutants and Omicron RBD

For further analysis, 6 single-point mutations for H11-H4 nanobody R27E, S54E, S57D, S107K, D108W, and A110I were shortlisted based on molecular docking scores and MD data **(Table 2)**. The interaction patterns at the interface of these mutants with RBD were identified through protein-protein docking using HADDOCK 2.4. and compared with native H11-H4 **(Figure7A)**. The RBD and H11-H4-R27E complex exhibited a total of six H-bonds with the RBD. Notably, the mutated residue E27 in CDR1 of H11-H4 formed two new H-bonds with R346 of the RBD. Additionally, S54 in CDR2 and H100 in CDR3 of H11-H4 formed new H-bonds with R493 and S496 of RBD, respectively **(Figure 7B)**. In the RBD: H11-H4-S54E complex the mutated residue E54 in CDR2 did not form any H-bond interactions. However, a framework region residue E44 formed an extra H-bond with K478 of RBD **(Figure 7C)**. For the RBD: H11-H4-S57D complex, a total of three H-bonds and a salt bridge were observed with RBD. The mutated residue D57 in CDR2 formed a salt bridge with R498 and an H-bond with Y449 of RBD. Additionally, residues A58 in CDR2 and K65 present in the framework region of H11-H4 formed a new H-bond with R498 and S446 of RBD **(Figure 7D)**. In the RBD: H11-H4-S107K complex, the mutated residue K107 in CDR3 did not form any new H-bond, but rather an extensive H-bond network was observed between CDR3 residues V102, P113, Y114, D115, and RBD residues F486, R493, and Y449 respectively. Additionally, framework region residues Q1, Y95, and, G118 formed H-bonds with Y449 and R498 of RBD respectively **(Figure 7E)**. For the RBD: H11-H4-D108W complex, a total of 6 H-bonds were observed with RBD. Mutated residue W108 in CDR3 formed a new H-bond with A484 of RBD. In addition to this RBD residue R498, S446, and Y501 were making new H-bonds with K74 in the framework region and T28 in CDR1 of the H11-H4 nanobody **(Figure 7F)**. For RBD: H11-H4-A110I complex, CDR3 residue H100 formed H-bond with Y449 and residues P113, Y114, and D115 were forming H-bonds with R493 residue of RBD. Additionally, framework residues G42, E44, and R45 formed H-bonds with RBD residues K478, G485, and F489 **(Figure 7G)**. This analysis revealed that mutations within specific CDR loops not only alter the binding characteristics at the mutation site but also influence the binding patterns in other CDRs and framework regions **(Figure 7 and Supplementary Figure 3)**. This finding highlights the intricate interplay between different regions of the nanobody and their collective role in determining overall binding efficiency. The analysis of the secondary structure of these mutants of nanobodies showed that overall, no major structural changes occurred after mutation in nanobodies **(Supplementary Table 8 and supplementary Figure 4)**.

**Figure 7:**
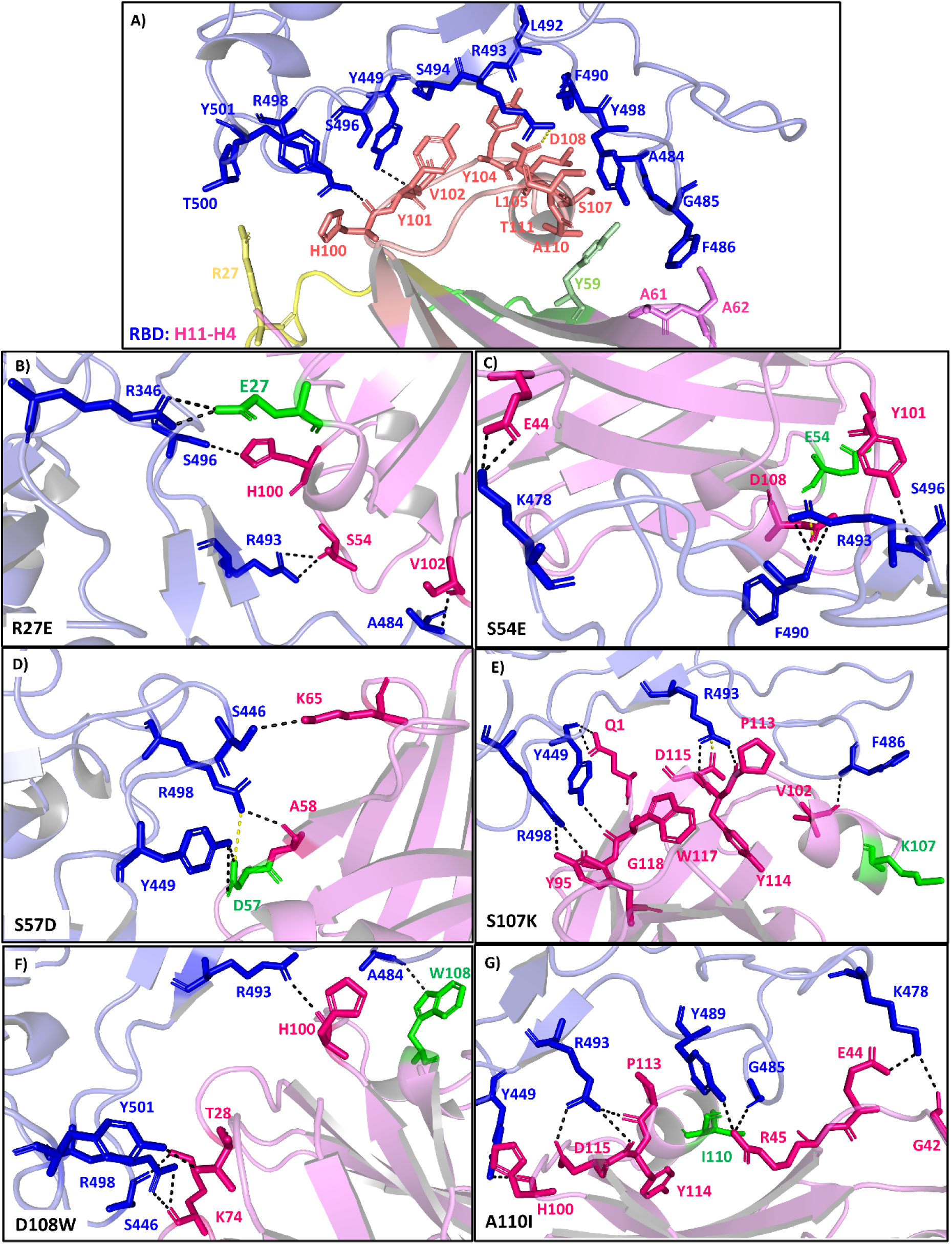
Interaction analysis of docked complex of Omicron RBD with H11-H4 nanobody and affinity matured mutants. A-G) Cartoon representation of Omicron RBD (slate) and H11-H4 (violet) nanobody complex showing H-bonded interaction present at the interface of Omicron RBD and nanobodies. A) H11-H4 nanobody CDR1 (yellow), CDR2 (green), and CDR3 (orange) residues interaction with RBD. B-G) Nanobody mutants R27E, S54E, S57D, S107K, D108W and A110I interactions with RBD. Residues of RBD participating in H-bonding are shown as blue sticks and labeled in blue text, H11-H4 nanobody residues are shown as hot pink sticks and labeled with hot pink text, and mutated residues of H11-H4 are shown as green sticks and labeled with green text. The dashed line indicates H-bonds (black) and Salt bridge (yellow).

**Table 2:** List of selected mutations of H11-H4 nanobody after *in silico* screening. Combined results for docking and MD analysis were given for each mutation.

| Mutation | HADDOCK 2.4 score | No. of H-bond/<br>docking | No. of Salt Bridge/<br>docking | No. of H-bond in<br>frame /MD | No. of Salt Bridge/MD | Average RMSD in<br>100 ns MD | No. of average<br>H-bond in 100<br>ns MD |
| --- | --- | --- | --- | --- | --- | --- | --- |
| **RBD+** | -81.3+/-17.2 | 2 | 1 | 6 | 1 | 0.61 | 9.5 |
| **H11-H4** |  |  |  |  |  |  |  |
| **R27E** | -91.4+/-3.0 | 6 | 0 | 14 | 0 | 0.36 | 6.3 |
| **S54E** | -85.3+/-10.4 | 6 | 0 | 15 | 1 | 0.59 | 11.1 |
| **S57D** | -84.8+/-4.0 | 3 | 1 | 13 | 0 | 0.57 | 9.6 |
| **S107K** | -87.7+/-11.5 | 5 | 1 | 11 | 1 | 0.48 | 10.7 |
| **D108W** | -88.9+/-1.5 | 6 | 0 | 9 | 0 | 0.61 | 6.3 |
| **A110I** | -85.6+/-12.0 | 7 | 0 | 11 | 0 | 0.49 | 10.5 |

In SARS-CoV-2 infection, the interaction between RBD and ACE2 receptor is crucial for viral entry into host cells (74). Specific mutations in nanobodies that disrupt this interaction can serve as potent inhibitors of viral entry. Analysis of the docked complexes showed that the RBD residues targeted by nanobodies were also engaged in binding with ACE2. Furthermore, certain nanobody mutations have been found to target additional residues within the ACE2 footprint of RBD **(Figure 8)**. Mutations in the H11-H4 nanobody, such as D108W, A110I, and R27E, were observed to interact with the Omicron RBD by occupying a larger surface area and engaging with more RBD residues **(Figures 8B, G, and, H)**. The ACE2 footprint is a prime target because of its essential role in viral entry, and antibodies that bind to this can effectively block the virus from initiating infection. These mutant nanobodies function as molecular roadblocks, obstructing the virus’s entry and subsequent infection. This dual targeting of critical residues by nanobodies suggests a promising therapeutic strategy against any future variants of SARS-CoV-2, offering a novel approach to block the virus at its initial point of contact with host cells.

**Figure 8:**
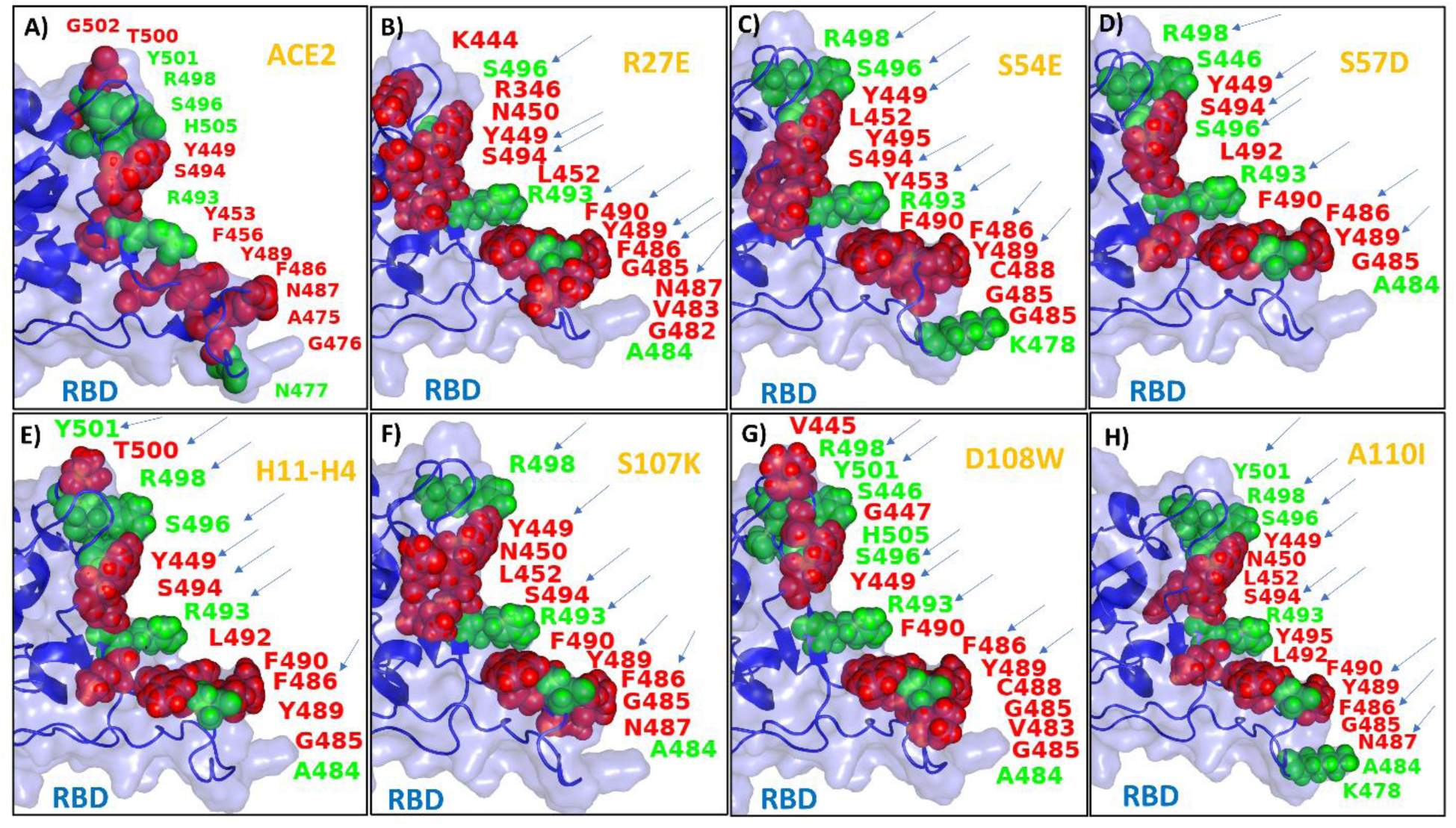
Omicron RBD highlighting the interacting residues shared between ACE2 and H11-H4 nanobody and their mutants. A-H) Omicron RBD is shown in surface (slate), cartoon (blue), and spheres represent the residue of RBD present within the 4Å cutoff of respective complex (Red: wild type; Green: mutated residue of Omicron). Arrows (blue) indicate the overlapping residues present at the interface of RBD: ACE2 and RBD: H11-H4 nanobody (native and its affinity matured mutations).

### 3.6 Biophysical assays substantiated the accuracy of the developed pipeline

Subtle structural modifications in the CDRs were likely to impact the binding interface between the Omicron RBD and nanobody, thereby modulating the binding affinities. To evaluate these interactions, Omicron RBD, H11-H4 nanobody, and selected mutants of H11-H4 were recombinantly produced and purified **(Supplementary Figure 5)**. ITC was used to measure the binding affinities and thermodynamic parameters. As anticipated, the interaction between the RBD and nanobody was primarily driven by exothermic processes, as indicated by the ΔH values **(Supplementary Table 9).** Thermodynamic data from ITC experiments were analyzed with a single-site binding model to evaluate and compare the binding affinities of the native nanobody (H11-H4) and its mutants. The H11-H4 nanobody shows a dissociation constant (K_D_) of ∼32 µM. Corroborating with *in silico* predictions, the single mutants D108W, S57D, R27E, A110I, and S107K exhibited enhanced binding affinities with K_D_ Values ∼8.8 µM, ∼10 µM, ∼14 µM, ∼15.6 µM and ∼27 µM respectively **(Figure 9)**. The significant binding affinity improvements in mutants D108W, S57D, R27E, and A110I corroborate *in silico* analysis, demonstrating that these targeted mutations enhance the interaction between the nanobody mutant and the RBD **(Figure 7)**. Such improvements underscore the potential of computational and experimental approaches in affinity maturation to develop effective therapeutic agents against SARS-CoV-2.

**Figure 9:**
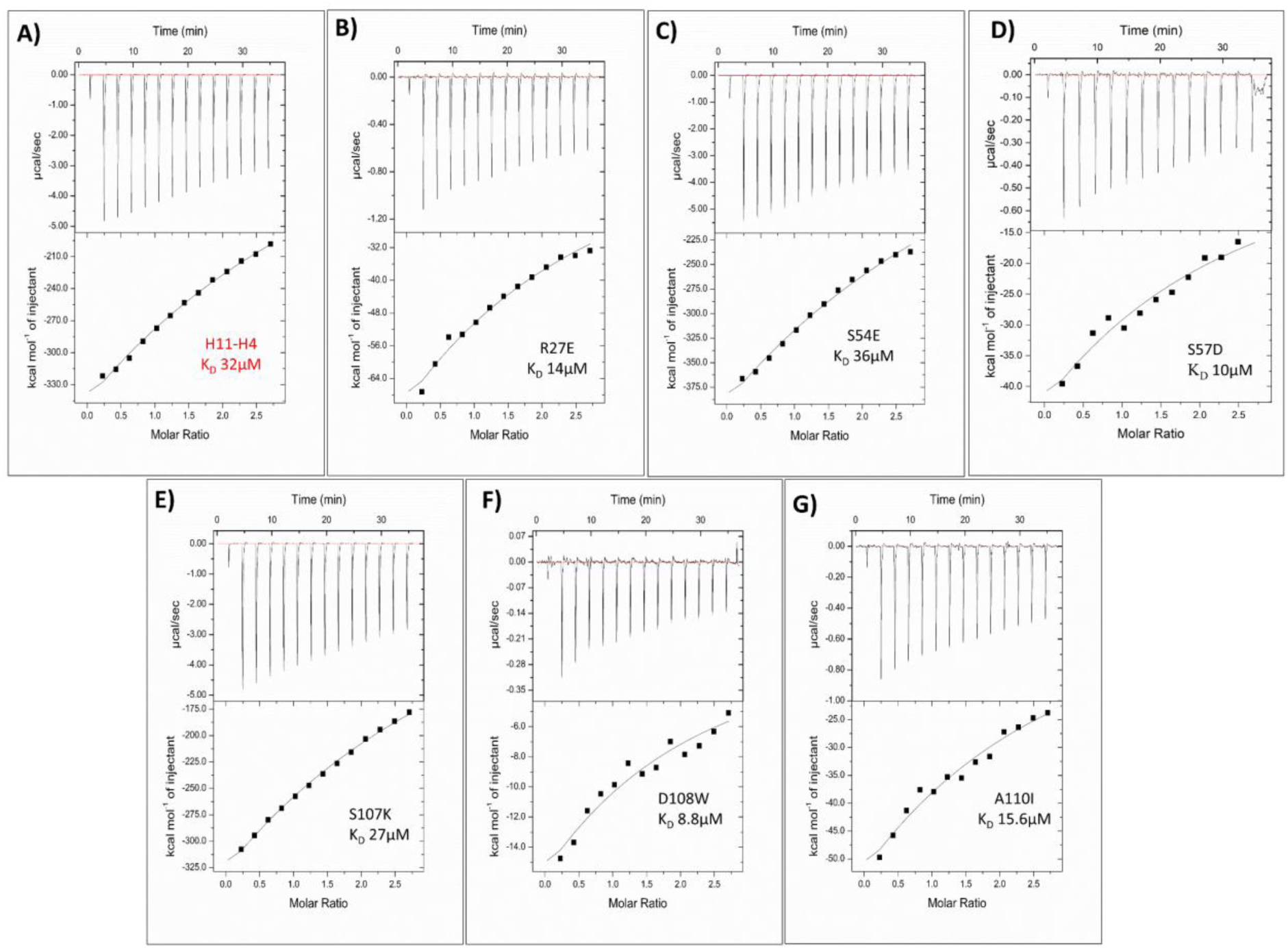
The binding affinity of native and mutant nanobodies towards the Omicron RBD was determined using ITC. Panels (A-G) display the binding isotherm plots, generated by applying a single-site binding model. The upper section of each plot shows the raw titration data, while the lower section presents the fitted binding isotherms. Inset K_D_ values for both native and mutant nanobodies are included for reference.

## 4. Discussion

Antibody-based therapies, such as monoclonal antibodies (mAbs) and nanobodies, provide an immediate and potent defense against SARS-CoV-2, especially crucial for immunocompromised individuals who may not respond effectively to vaccines. Unlike vaccines, which rely on stimulating the body’s immune response to produce antibodies, these therapies supply the antibodies directly to patients, offering immediate protection or treatment (75). This approach is especially valuable in addressing the challenges posed by rapid viral evolution and the emergence of new variants, as seen with SARS-CoV-2 (38,76,77). Nanobodies offer a rapid and robust alternative to conventional mAbs and represent a promising alternative for targeted prophylactic and therapeutic applications. ALX-0171, a therapeutic nanobody administered via nebulization, is currently undergoing clinical trials against respiratory syncytial virus (RSV) (78,79). Nanobodies can be supplemented through infant milk formulation to treat Group A Rotavirus (RVA) diarrhea (80).

The S-protein of the SARS-CoV-2, a primary immunogenic region, is targeted by neutralizing antibodies, Fabs, scFvs, and nanobodies. In this study, nanobodies H11-H4, H3, and C5, initially identified against RBD of Wuhan isolate of SARS-CoV-2, were repurposed to specifically target the Omicron RBD. All three nanobodies were docked with the Omicron RBD, and *in silico* analysis of the Omicron RBD and nanobody complexes revealed reduced interactions at the complex interface due to E484A, Q493R, and N501Y mutations **(Figure 1 and Supplementary Figures 1)**. The Omicron RBD exhibits 15 mutations, including N501Y, S477N, Q493R, and T478K, which enhance binding affinity towards ACE2, and other mutations like K417N, Q493R, G446S, and E484A, which enable immune evasion, rendering many established neutralizing antibodies ineffective (62,64,71–73). In antibody-antigen complex, CDRs play a pivotal role in antigen binding, contributing approximately 80% of the binding energy (81). These serve as mutational hotspots, influencing the binding specificity and affinity of nanobodies (13,23,82). Herein, Omicron RBD and nanobody complexes were subjected to a streamlined *in silico* affinity maturation pipeline to identify high affinity and stable binders. High-throughput *in silico* mutagenesis was employed to expand paratope diversity in nanobodies targeting the Omicron RBD. A comprehensive library of mutants was generated for each nanobody, mimicking the natural somatic hypermutation process to improve affinity and expand binding specificity. The docked complexes of Omicron RBD and nanobody (H11-H4, C5, and H3) undergo the virtual mutagenesis combining mCSM-AB2, DUET, and Deep DDG software resulting in an *in-house* library of ∼1,976 potentially high affinity and stable substitution mutations in the CDRs of selected nanobodies. This comprehensive approach identified mutations that enhance both binding affinity and stability **(Figure 2)**. A heat map illustrating ΔΔG contributions to binding affinity showed a preference for aromatic and charged residues, while ΔΔG contributions to stability favored non- polar and aromatic residues **(Figure 3)**. These findings are consistent with the general findings of alanine-scanning mutagenesis studies, which indicate that only a few side chains predominantly contribute to the energy of association in a functional epitope, typically those that are charged or aromatic (70,81,83,84).

Following extensive virtual mutagenesis in CDRs of nanobodies, 35 high-affinity, stable mutations were identified for H11-H4 and H3 nanobodies, and 26 for C5 nanobody, predominantly involving aromatic or charged residues mutations **(Figure 4 and Supplementary Tables 1, 3, and 5)**. These mutants were subsequently docked with the Omicron RBD, and mutations that contributed to more favorable interactions were selected 18 mutants for H11-H4, 14 mutants for H3, and 8 mutants for C5 **(Supplementary Tables 1, 3, and 5)**. Further screening and selection of these mutants were based on the MD simulation studies, providing insights into the structural integrity and stability of the Omicron RBD and nanobody complex after mutation **(Figure 4, Supplementary Figure 2 and Supplementary Table 2, 4, and 6)**. Analysis of the mutational landscape of the H11-H4 nanobody emphasized the importance of surface complementarity. As per the examination, specific amino acid positioning in CDRs was identified as a critical factor affecting binding affinity with the RBD **(Figures 6A and 6B)**. Investigation of Omicron RBD interactions with H11-H4 nanobody and ACE2 reveals distinct residue distributions. Aromatic and charged residues dominated both interfaces, crucial for binding affinity. Mutational trends in nanobody affinity maturation emphasize enhancing aromatic and charged interactions with RBD **(Figures 4, 6B, and 6C).** The current methodology identified six unique and stable binders for the H11-H4 nanobody R27E, S54E, S57D, S107K, D108W, and A110I. For the C5 and H3 nanobodies, the selected high-affinity binders were H32Y, I57E, G99V, G99I, and, Y109E and R27E, T54D, T100E, T107D, G115V and, S116I respectively **(Supplementary Tables 2, 4, and 6)**. During affinity maturation, antibodies often accumulate charged mutations in their CDRs to enhance electrostatic interactions (84). This phenomenon is crucial as the binding interfaces of antibody-antigen complexes often comprise a substantial proportion of aromatic and charged residues, contrasting with protease-inhibitor complexes where such residues are less prevalent (70,81,83). The identified mutations in the affinity-matured nanobodies align with these observations, underscoring their pivotal role in augmenting affinity for the Omicron RBD.

Detailed analyses of docked complexes of Omicron RBD and mutant nanobodies reveal that mutations in specific CDRs affect binding characteristics both within those CDRs and across other CDRs and the framework region, highlighting the complex interplay that determines binding efficiency. **(Figure 5 and Supplementary Figures 3)**. The nanobody mutants D108W, S57D, R27E, and A110I formed a more extensive network of hydrogen and hydrophobic bonds with the Omicron RBD **(Figure 7)**. Mutations in the H11-H4 nanobody, such as D108W, A110I, and R27E, interacted with the Omicron RBD by covering a larger surface area and engaging more RBD residues causing steric hindrance for ACE2 binding **(Figure 8).** To validate the *in silico* findings, the H11-H4 and its mutant nanobodies were produced recombinantly, and their binding affinities with Omicron RBD were evaluated *in vitro* using ITC **(Supplementary Table 9)**. The mutants R27E, S57D, S107K, D108W, and A110I have K_D_ Values ranging from ∼8.8 to ∼ 27 µM which were significantly better than native nanobody (∼32 µM) **(Figure 9).** This study successfully repurposed the H11-H4 nanobody through an *in silico* affinity maturation pipeline, achieving approximately three times better affinity than the native H11-H4 nanobody by substituting single amino acids.

In summary, leveraging structure-guided redesigning offers a rapid method to engineer existing neutralizing nanobodies or antibodies for new applications, akin to drug repurposing in drug discovery. The developed protocol is both time and cost-efficient, facilitating the development of prophylactic immunization against various viral pathogens, including SARS-CoV-2. To advance these mutant nanobodies as therapeutic agents, humanization and optimization processes are crucial to enhance their avidity and efficacy (46,85–87). This study provides preliminary data, and further validation through *in vitro*, and *in vivo* characterization studies cannot be ruled out and is warranted to avail the benefits of these small-size biologics.

## 5. Conclusion

This study successfully repurposed the nanobodies H11-H4, H3, and C5, initially developed against the Wuhan RBD, to specifically target the Omicron RBD of SARS-CoV-2. A robust structure guided *in silico* affinity maturation pipeline was used to introduce strategic mutations in the CDRs to enhance the binding affinity of nanobodies towards Omicron RBD. This pipeline integrates high-throughput mutagenesis, protein-protein docking, and MD simulations, to shortlist high affinity mutants from an extensive *in house* mutant library. Experimental validation using ITC confirmed that the affinity-matured H11-H4 mutants, augmented with aromatic and charged residues, exhibited significantly improved binding with omicron RBD, having up to a three-fold increase in binding affinity compared to the native nanobody. This approach offers a rapid and cost efficient strategy to bioengineer existing nanobodies to counter emerging viral variants.

## Funding

The Indian Council for Medical Research (ICMR) Government of India (Project. ref no ISRM/12/ (06)/2022) supported this study.

## CRediT authorship contribution statement

**Vishakha Singh**: Conceptualization, Data curation, Formal analysis, Software, Methodology, Validation, Visualization, Writing – original draft. **Shweta Choudhary**: Methodology, Visualization, Validation Writing – original draft. **Mandar Bhutkar**: Methodology, Writing – review & editing. **Sanketkumar Nehul**: Methodology, Validation. **Sabika Ali**: Methodology. **Jitin Singla**: Conceptualization, Formal analysis, Software, Supervision, Validation, Writing – review & editing, Funding acquisition. **Pravindra Kumar**: Supervision, Resources, Validation, Writing – review & editing. **Shailly Tomar**: Conceptualization, Formal analysis, Funding acquisition, Methodology, Validation, Project administration, Resources, Supervision, Writing – review & editing.

## Acknowledgment

ST and JS acknowledge and thank the Indian Council for Medical Research (ICMR) Government of India (Project. ref no ISRM/12/ (06)/2022) for supporting this study. VS is thankful to ICMR, SC, and MB are thankful to the Council of Scientific and Industrial Research (CSIR) for financial support. SKN and SA acknowledge the Ministry of Human Resource Development, (MHRD) for research fellowship. ST, JS, and PK thank the Department of Biotechnology, Govt of India for supporting the Translational and Structural Bioinformatics Centre at the Department of Biosciences and Bioengineering, IIT Roorkee (reference number BT/PR40141/BTIS/137/16/2021). The authors also thank Ashok Soota Molecular Medicine Facility and Macromolecular Crystallographic Unit (MCU) at the Indian Institute of Technology Roorkee (IIT Roorkee).

## Declaration of competing interest

We declare that we have no conflicts of interest in the authorship or publication of this contribution.

## Data availability

Data will be made available upon reasonable request to the corresponding author.

## Supplementary information

### Supplementary Tables

**Supplementary Table 1:**
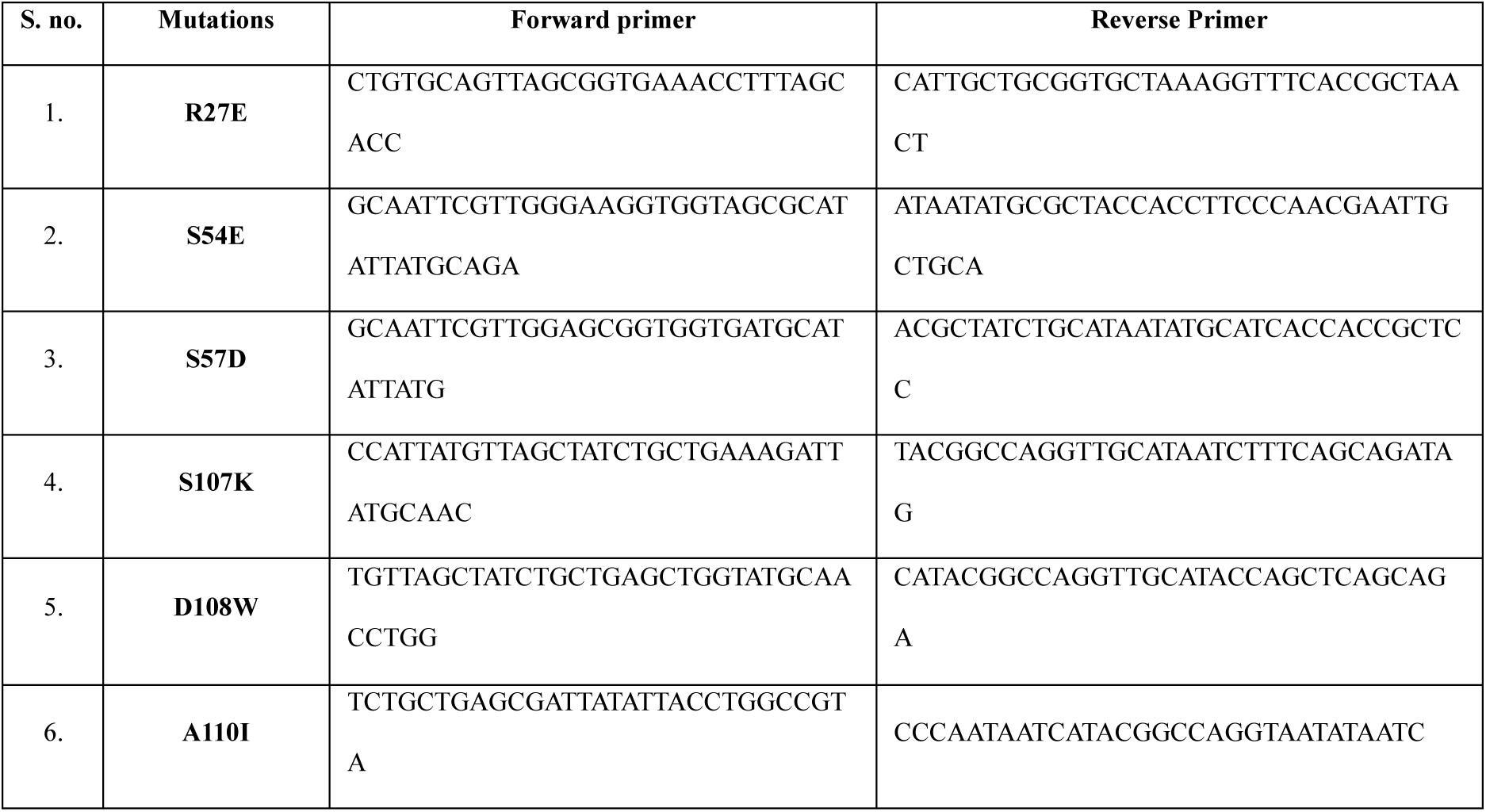
Set of primers used for the Site Directed Mutagenesis of H11-H4 nanobody.

**Supplementary Table 2:** Selected mutations for H11-H4 nanobody after *in silico* affinity maturation using mCSMAb2, DUET, and Deep DDG softwares, along with their docking scores and interactions analysis. Mutations selected post-docking for the H11-H4 nanobody are highlighted in green, while the docking results for the native nanobody are shown in red.

| S.no | Wild | Position | Mutant | DDG<br>(mCSMAb2) | ddG (kcal/mol,<br>>0 is stable, <0 is<br>unstable) | Haddock score | H-bond | Salt Bridge |
| --- | --- | --- | --- | --- | --- | --- | --- | --- |
|  |  |  |  |  |  | -81.3+/-17.2 | 2 | 1 |
| 1 | R | 27 | E | 0.22 | 0.461 | -91.4+/-3.0 | 6 | 0 |
| 2 | R | 27 | F | 0.96 | 0.301 | -94.7+/-2.9 | 2 | 0 |
| 3 | R | 27 | Y | 0.55 | 0.137 | -88.0+/-5.1 | 3 | 0 |
| 4 | A | 33 | L | 0.11 | 0.424 | -87.4+/-3.2 | 3 | 0 |
| 5 | A | 50 | V | 0.96 | 0.195 | -87.6+/-8.7 | 7 | 0 |
| 6 | A | 50 | L | 1.3 | 0.293 | -84.5+/-10.7 | 6 | 0 |
| 7 | A | 50 | I | 1.47 | 0.407 | -89.6+/-10.7 | 3 | 0 |
| 8 | S | 54 | D | 0.3 | 0.378 | -88.1+/-13.9 | 7 | 0 |
| 9 | S | 54 | E | 0.24 | 0.246 | -85.3+/-10.4 | 6 | 0 |
| 10 | S | 57 | D | 0.18 | 0.120 | -84.8+/-4.0 | 3 | 1 |
| 11 | S | 57 | E | 0.2 | 0.168 | -84.9+/-1.4 | 5 | 0 |
| 12 | A | 58 | V | 0.55 | 0.230 | -87.9+/-10.7 | 8 | 1 |
| 13 | A | 58 | L | 0.12 | 0.169 | -84.9+/-5.3 | 3 | 0 |
| 14 | A | 58 | I | 0.88 | 0.273 | -73.7 +/- 9.8 | 3 | 0 |
| 15 | H | 100 | L | 0.19 | 0.407 | -72.9+/-2.7 | 3 | 0 |
| 16 | H | 100 | W | 0.62 | 0.491 | -84.8+/-10.7 | 4 | 1 |
| 17 | H | 100 | I | 0.16 | 0.269 | -93.0+/-6.3 | 6 | 0 |
| 18 | H | 100 | F | 0.15 | 0.626 | -80.3+/-4.2 | 2 | 0 |
| 19 | Y | 101 | W | 0.73 | 0.358 | -80.5+/-9.5 | 2 | 1 |
| 20 | S | 103 | T | 0.24 | 0.167 | -90.6+/-18.4 | 8 | 0 |
| 21 | S | 103 | K | 0.76 | 0.046 | -84.9+/-1.7 | 2 | 0 |
| 22 | S | 107 | K | 0.06 | 0.158 | -87.7+/-11.5 | 5 | 1 |
| 23 | D | 108 | E | 0.15 | 0.051 | -82.6+/-2.2 | 2 | 0 |
| 24 | D | 108 | F | 0.22 | 0.275 | -101.8+/-1.5 | 10 | 0 |
| 25 | D | 108 | Y | 1.39 | 0.300 | -88.9+/-1.5 | 10 | 0 |
| 26 | D | 108 | W | 3.34 | 0.091 | -88.9+/-1.5 | 6 | 0 |
| 27 | A | 110 | V | 0.45 | 0.585 | -82.2+/-7.7 | 6 | 0 |
| 28 | A | 110 | L | 1.23 | 1.124 | -82.4+/-8.5 | 6 | 0 |
| 29 | A | 110 | M | 0.74 | 0.558 | -83.8+/-3.2 | 4 | 0 |
| 30 | A | 110 | I | 1.1 | 0.718 | -85.6+/-12.0 | 7 | 0 |
| 31 | A | 110 | C | 0.56 | 0.053 | -76.5+/-3.6 | 3 | 0 |
| 32 | A | 110 | P | 0.55 | 0.335 | -93.0+/-12.3 | 6 | 0 |
| 33 | A | 110 | E | 3.36 | 0.007 | -80.2+/-8.4 | 8 | 0 |
| 34 | T | 111 | E | 0.71 | 0.147 | -83.6+/-8.4 | 2 | 0 |
| 35 | T | 111 | D | 0.71 | 0.038 | -89.6+/-2.1 | 2 | 0 |

**Supplementary Table 3:**
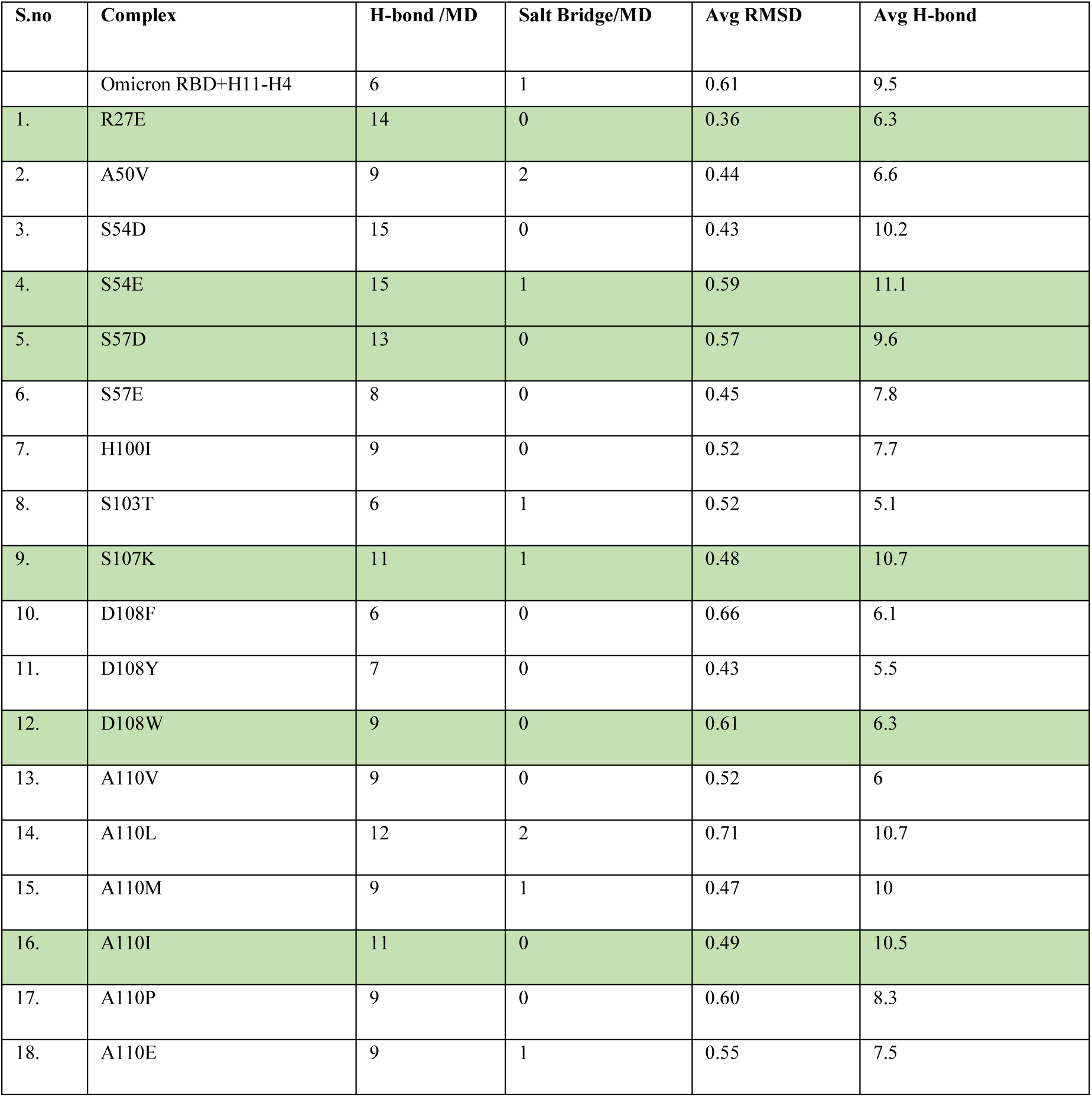
List of mutations for H11-H4 nanobody displaying increased number of interactions with Omicron-RBD than native after MD. The frame PBDs were extracted from the MD run trajectories and used for the generation of DimPlots. Mutations selected after MD simulations were highlighted in green.

| S.no | Complex | H-bond /MD | Salt Bridge/MD | Avg RMSD | Avg H-bond |
| --- | --- | --- | --- | --- | --- |
|  | Omicron RBD+H11-H4 | 6 | 1 | 0.61 | 9.5 |
| 1. | R27E | 14 | 0 | 0.36 | 6.3 |
| 2. | A50V | 9 | 2 | 0.44 | 6.6 |
| 3. | S54D | 15 | 0 | 0.43 | 10.2 |
| 4. | S54E | 15 | 1 | 0.59 | 11.1 |
| 5. | S57D | 13 | 0 | 0.57 | 9.6 |
| 6. | S57E | 8 | 0 | 0.45 | 7.8 |
| 7. | H100I | 9 | 0 | 0.52 | 7.7 |
| 8. | S103T | 6 | 1 | 0.52 | 5.1 |
| 9. | S107K | 11 | 1 | 0.48 | 10.7 |
| 10. | D108F | 6 | 0 | 0.66 | 6.1 |
| 11. | D108Y | 7 | 0 | 0.43 | 5.5 |
| 12. | D108W | 9 | 0 | 0.61 | 6.3 |
| 13. | A110V | 9 | 0 | 0.52 | 6 |
| 14. | A110L | 12 | 2 | 0.71 | 10.7 |
| 15. | A110M | 9 | 1 | 0.47 | 10 |
| 16. | A110I | 11 | 0 | 0.49 | 10.5 |
| 17. | A110P | 9 | 0 | 0.60 | 8.3 |
| 18. | A110E | 9 | 1 | 0.55 | 7.5 |

**Supplementary Table 4:** Selected mutations for C5 nanobody after *in silico* affinity maturation using mCSMAb2, DUET, and Deep DDG softwares, along with their docking scores and interactions analysis. Mutations selected post-docking for the C5 nanobody are highlighted in green, while the docking results for the native nanobody are shown in red.

| S.no | Wild | Position | Mutant | DDG (mCSM-AB2) | ddG (kcal/mol, >0 is stable, <0 is unstable) | HADDOCK score | H-bond/ docking |
| --- | --- | --- | --- | --- | --- | --- | --- |
|  |  |  |  |  |  | -103+/-5.6 | 5 |
| 1 | H | 32 | Y | 2.48 | 0.814 | -105.8+/-0.5 | 8 |
| 2 | H | 32 | F | 0.48 | 0.325 | -78.4+/-12.4 | 3 |
| 3 | H | 32 | W | 0.41 | 0.499 | -122.7+/-3.8 | 4 |
| 4 | I | 57 | K | 0.07 | 0.105 | -126.3+/-2.7 | 5 |
| 5 | I | 57 | R | 0.33 | 0.256 | -125+/-2.8 | 5 |
| 6 | I | 57 | H | 0.46 | 0.001 | -127.7+/-1.5 | 3 |
| 7 | I | 57 | D | 0.29 | 0.116 | -123.4+/-1.3 | 6 |
| 8 | I | 57 | E | 0.27 | 0.247 | -128.6+/-1.1 | 6 |
| 9 | I | 57 | N | 0.36 | 0.137 | -124.4+/-1.6 | 5 |
| 10 | G | 99 | V | 2.03 | 0.262 | -107.0+/-4.5 | 6 |
| 11 | G | 99 | L | 2.2 | 0.723 | -123.9+/-1.6 | 4 |
| 12 | G | 99 | I | 2.15 | 0.513 | -112.3+/-8.5 | 6 |
| 13 | A | 103 | G | 0.23 | 0.335 | -126.6+/-0.8 | 3 |
| 14 | A | 103 | V | 0.18 | 0.071 | -126+/-1.7 | 5 |
| 15 | A | 103 | R | 0.38 | 0.100 | -124.5+/-1.2 | 5 |
| 16 | A | 103 | S | 0.13 | 0.220 | -126.7+/-2.9 | 3 |
| 17 | S | 105 | N | 0.26 | 0.046 | -125.5+/-3.2 | 5 |
| 18 | D | 106 | E | 0.21 | 0.117 | -128.8+/-1.8 | 5 |
| 19 | N | 107 | Q | 0 | 0.023 | -127.4+/-1.8 | 5 |
| 20 | N | 107 | S | 0.08 | 0.199 | -124.2+/-1.2 | 6 |
| 21 | N | 107 | T | 0 | 0.423 | -122.9+/-0.9 | 6 |
| 22 | N | 107 | D | 0.2 | 0.186 | -125.6+/-1.6 | 3 |
| 23 | N | 107 | E | 0.06 | 0.454 | -123.3+/-3.3 | 4 |
| 24 | Y | 109 | D | 0.19 | 0.109 | -124.1+/-0.9 | 5 |
| 25 | Y | 109 | E | 0.05 | 0.176 | -126.2+/-1.7 | 7 |
| 26 | F | 110 | Y | 1.26 | 0.204 | -120.7+/-2.4 | 4 |

**Supplementary Table 5:** List of mutations for C5 nanobody displaying increased number of interactions with Omicron-RBD than native after MD. The frame PBDs were extracted from the MD run trajectories and used for the generation of DimPlots. Mutations selected after MD simulations were highlighted in green.

| S.no | Complex | No. of H- bond / frame | Avg RMSD | Avg H- bond |
| --- | --- | --- | --- | --- |
|  | Omicron RBD+C5 | 10 | 0.42 | 7.4 |
| 1. | H32Y | 15 | 0.51 | 8.85 |
| 2. | I57D | 7 | 0.43 | 8.2 |
| 3. | I57E | 14 | 0.55 | 8.16 |
| 4. | G99V | 15 | 0.55 | 9.35 |
| 5. | G99I | 16 | 0.47 | 8.64 |
| 6. | N107S | 7 | 0.6 | 7.83 |
| 7. | N107T | 6 | 0.75 | 9.51 |
| 8. | Y109E | 12 | 0.5 | 7.44 |

**Supplementary Table 6:** Selected mutations for H3 nanobody after *in silico* affinity maturation using mCSMAb2, DUET, and Deep DDG softwares, along with their docking scores and interactions analysis. Mutations selected post-docking for the H3 nanobody are highlighted in green, while the docking results for the native nanobody are shown in red.

| S.no | Wild | Position | Mutant | DDG (mCSMAb2) | ddG (kcal/mol, >0 is stable, <0 is unstable) | Haddock score | H- bond | Salt Bridge |
| --- | --- | --- | --- | --- | --- | --- | --- | --- |
|  |  |  |  |  |  | -90.1+/-2.5 | 3 | 0 |
| 1 | R | 27 | V | 0.01 | 0.238 | -89.2+/-9.6 | 5 | 0 |
| 2 | R | 27 | T | 0.04 | 0.161 | -80.0+/-1.0 | 7 | 0 |
| 3 | R | 27 | N | 0.02 | 0.052 | -90.2+/-2.4 | 6 | 0 |
| 4 | R | 27 | Q | 0.02 | 0.052 | -90.0+/-7.9 | 5 | 0 |
| 5 | R | 27 | D | 0.21 | 0.154 | -92.9+/-3.7 | 5 | 0 |
| 6 | R | 27 | E | 0.26 | 0.142 | -86.8+/-7.1 | 5 | 1 |
| 7 | R | 27 | F | 0.53 | 0.018 | -88.9+/-5.8 | 5 | 0 |
| 8 | T | 28 | E | 0.19 | 0.098 | -95.3+/-2.5 | 6 | 0 |
| 9 | T | 31 | E | 0.08 | 0.366 | -91.4+/-4.0 | 3 | 0 |
| 10 | T | 31 | D | 0.25 | 0.321 | -90.4+/-4.0 | 2 | 0 |
| 11 | M | 51 | I | 2.33 | 0.513 | -93.0+/-1.1 | 5 | 0 |
| 12 | T | 54 | D | 0.03 | 0.329 | -87.6+/-3.1 | 5 | 1 |
| 13 | S | 56 | E | 0.25 | 0.188 | -107.2+/-3.8 | 6 | 0 |
| 14 | S | 56 | N | 0.45 | 0.130 | -85.5+/-4.1 | 3 | 0 |
| 15 | T | 100 | E | 0.5 | 0.123 | -97.9+/-6.1 | 7 | 0 |
| 16 | T | 100 | D | 0.28 | 0.149 | -89.7+/-1.9 | 3 | 0 |
| 17 | V | 102 | R | 0.8 | 0.046 | -90.7+/-2.4 | 2 | 0 |
| 18 | A | 104 | L | 1.29 | 0.294 | -81.2+/-2.8 | 6 | 0 |
| 19 | A | 104 | P | 0.31 | 0.036 | -86.7+/-5.6 | 5 |  |
| 20 | Y | 105 | W | 1.25 | 0.037 | -87.0+/-8.7 | 6 | 0 |
| 21 | Y | 106 | F | 0.11 | 0.108 | -96.6+/-3.8 | 5 | 0 |
| 22 | T | 107 | E | 0.41 | 0.616 | -103.9+/-8.2 | 6 | 0 |
| 23 | T | 107 | D | 0.4 | 0.394 | -103.0+/-5.3 | 9 | 0 |
| 24 | Y | 109 | W | 0.18 | 0.507 | -87.0+/-8.7 | 6 | 0 |
| 25 | D | 113 | W | 0.08 | 0.044 | -94.5+/-2.9 | 6 | 0 |
| 26 | G | 115 | D | 0.54 | 0.368 | -88.5+/-4.3 | 3 | 0 |
| 27 | G | 115 | V | 0.01 | 0.650 | -87.7 +/- 1.5 | 7 | 0 |
| 28 | G | 115 | M | 0.78 | 0.182 | -83.7+/-3.7 | 4 | 0 |
| 29 | G | 115 | T | 0.09 | 0.144 | -89.5+/-4.2 | 4 | 0 |
| 30 | G | 115 | C | 0.14 | 0.073 | -93.6+/-2.7 | 3 | 1 |
| 31 | G | 115 | E | 0.82 | 0.458 | -79.3+/-0.7 | 3 | 0 |
| 32 | S | 116 | L | 0.11 | 0.083 | -94.4+/-4.7 | 7 | 0 |
| 33 | S | 116 | D | 0.14 | 0.197 | -91.2+/-0.6 | 5 | 0 |
| 34 | S | 116 | I | 0 | 0.035 | -88.0+/-3.1 | 9 | 1 |
| 35 | S | 116 | E | 0.07 | 0.111 | -83.8+/-2.0 | 5 | 0 |

**Supplementary Table 7:** List of mutations for H3 nanobody displaying increased number of interactions with Omicron-RBD than native after MD. The frame PBDs were extracted from the MD run trajectories and used for the generation of DimPlots. Supplementary Table 5: List of mutations for C5 nanobody displaying increased number of interactions with Omicron-RBD than native after MD. The frame PBDs were extracted from the MD run trajectories and used for the generation of DimPlots. Mutations selected after MD simulations were highlighted in green.

| S.no | Complex | H-bond | Salt Bridge | Avg RMSD | Avg H-bond |
| --- | --- | --- | --- | --- | --- |
|  | Omicron RBD+H3 | 7 | 0 | 0.5 | 7.71 |
| 1. | R27E | 1 | 0 | 0.69 | 3.22 |
| 2. | R27T | 4 | 0 | 0.55 | 5.89 |
| 3. | R27N | 11 | 1 | 0.47 | 8.61 |
| 4. | T28E | 6 | 0 | 0.55 | 5.56 |
| 5. | T54D | 14 | 0 | 0.5 | 10.39 |
| 6. | S56E | 8 | 1 | 0.5 | 5.59 |
| 7. | T100E | 12 | 0 | 0.59 | 7.55 |
| 8. | A104L | 3 | 0 | 0.77 | 3.45 |
| 9. | Y105W | 7 | 0 | 0.35 | 6.04 |
| 10. | T107D | 11 | 0 | 0.51 | 6.5 |
| 11. | D113W | 7 | 0 | 0.52 | 5.23 |
| 12. | G115V | 16 | 0 | 0.9 | 8.5 |
| 13. | S116I | 13 | 0 | 0.56 | 8.25 |
| 14. | S116L | 6 | 0 | 0.52 | 6.82 |

**Supplementary Table 8:** The secondary structure predictions of the native nanobody and its mutants was performed with DSSP over the 100ns MD simulations.

| S.No | Nanobody | Structure* | Coil | B-Sheet | B-Bridge | Bend | Turn | A-Helix | 3-Helix |
| --- | --- | --- | --- | --- | --- | --- | --- | --- | --- |
| 1. | Native (H11-H4) | 0.58 | 0.27 | 0.46 | 0.01 | 0.13 | 0.09 | 0.02 | 0.02 |
| 2. | R27E | 0.62 | 0.25 | 0.48 | 0.01 | 0.11 | 0.09 | 0.04 | 0.02 |
| 3. | S54E | 0.59 | 0.28 | 0.46 | 0.02 | 0.12 | 0.07 | 0.03 | 0.01 |
| 4. | S57D | 0.60 | 0.26 | 0.47 | 0.01 | 0.12 | 0.10 | 0.02 | 0.02 |
| 5. | S107K | 0.57 | 0.25 | 0.48 | 0.01 | 0.18 | 0.04 | 0.04 | 0.00 |
| 6. | D108W | 0.60 | 0.26 | 0.47 | 0.01 | 0.12 | 0.10 | 0.03 | 0.02 |
| 7. | A110I | 0.57 | 0.26 | 0.46 | 0.01 | 0.13 | 0.10 | 0.00 | 0.04 |

**\*Structure** = A-Helix + B-Sheet + B-Bridge + Turn

**Supplementary Table 9:** The thermodynamic analysis of the RBD and H11-H4 and its mutant as obtained from ITC.

| S.no. | Name | No. of sites (N) | K<sub>D</sub> (μM) | Δ H (cal/mol) | ΔS(cal/mol/degree) |
| --- | --- | --- | --- | --- | --- |
| 1. | **H11-H4(Native)** | 1 | 32μM | (-4.72)10<sup>6</sup>± (1.54)10<sup>5</sup> | (-1.58)10<sup>4</sup> |
| 2. | **H11-H4+R27E** | 1 | 14μM | (-4.56) 10<sup>5</sup>± (2.44)10<sup>4</sup> | (-1.51)10<sup>3</sup> |
| 3. | **H11-H4+S54E** | 1 | 36μM | (-5.93) 10<sup>6</sup>± (2.68) 10<sup>5</sup> | (-1.99) 10<sup>4</sup> |
| 4. | **H11-H4+S57D** | 1 | 10 μM | (-2.12) 10<sup>5</sup>± (1.86)10<sup>4</sup> | -690 |
| 5. | **H11-H4+S107K** | 1 | 27μM | (-3.78) 10<sup>6</sup>± (8.17)10<sup>4</sup> | (-1.27)10<sup>4</sup> |
| 6. | **H11-H4+D108W** | 1 | 8.8μM | (-6.78) 10<sup>4</sup>± 6606 | -204 |
| 7. | **H11-H4+A110I** | 1 | 15.6μM | (-3.62)10<sup>5</sup>± (2.68)10<sup>4</sup> | (-1.19)10<sup>3</sup> |

### Supplementary Figures

**Supplementary Figure 1:**
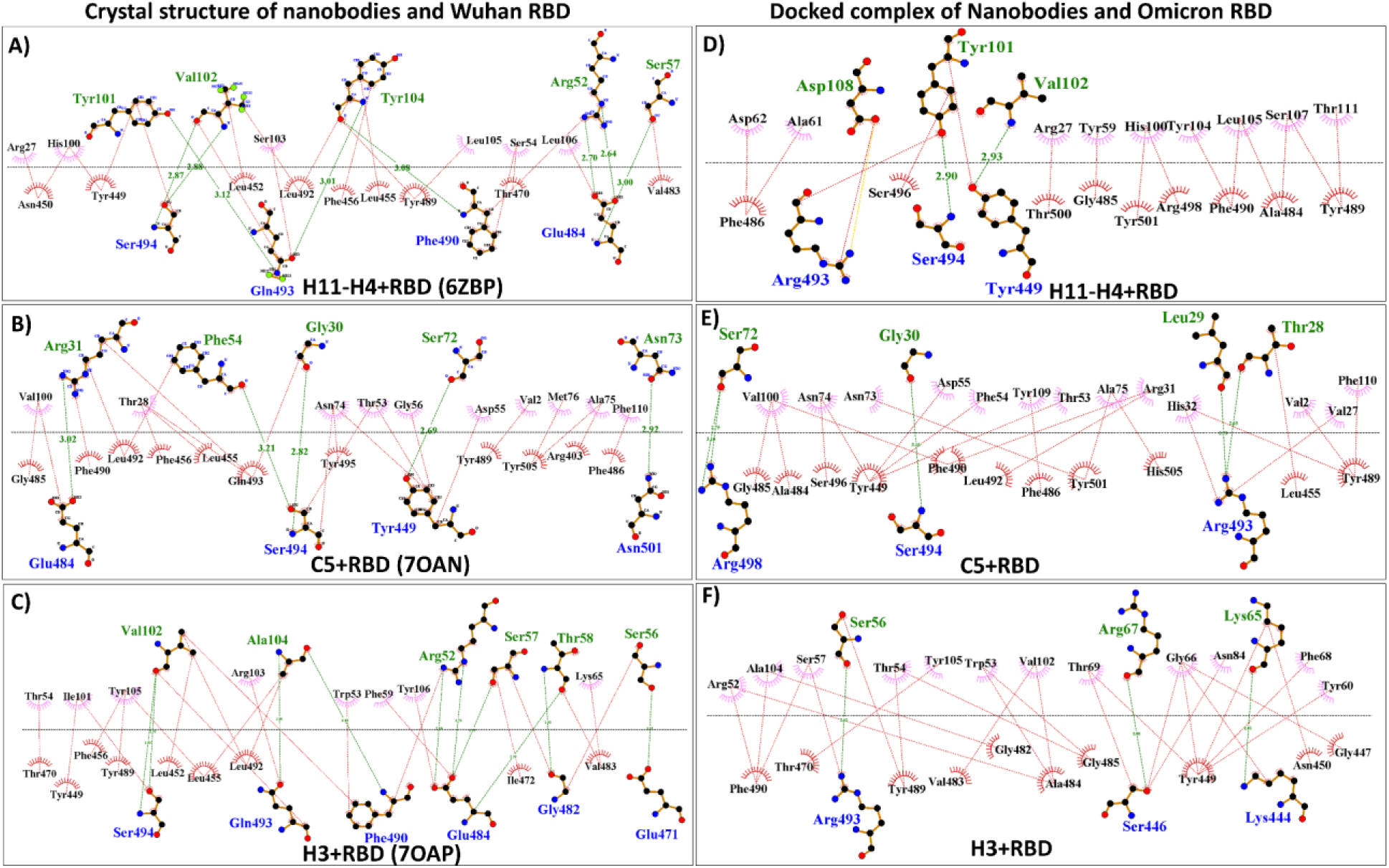
Comparative analysis showing 2D schematic representation of nanobody and RBD interacting residues generated using Dimplot. A-C). Showing interactions between Wuhan-RBD and nanobody using PDB IDs: 6ZBP (H11-H4), 7OAN (C5), and 7OAP (H3). D-F). Showing interactions between Omicron RBD and nanobody using complex generated through HADDOCK 2.4. RBD residues were shown in green colour and nanobody residue were shown in blue colour. Hydrogen bonds are depicted by green dashed lines connecting the atoms involved, with the donor-acceptor distance indicated in Å. Salt bridges are depicted as yellow dashed lines and hydrophobic contacts are shown as red arcs with spokes extending towards the atoms they interact with.

**Supplementary Figure 2:**
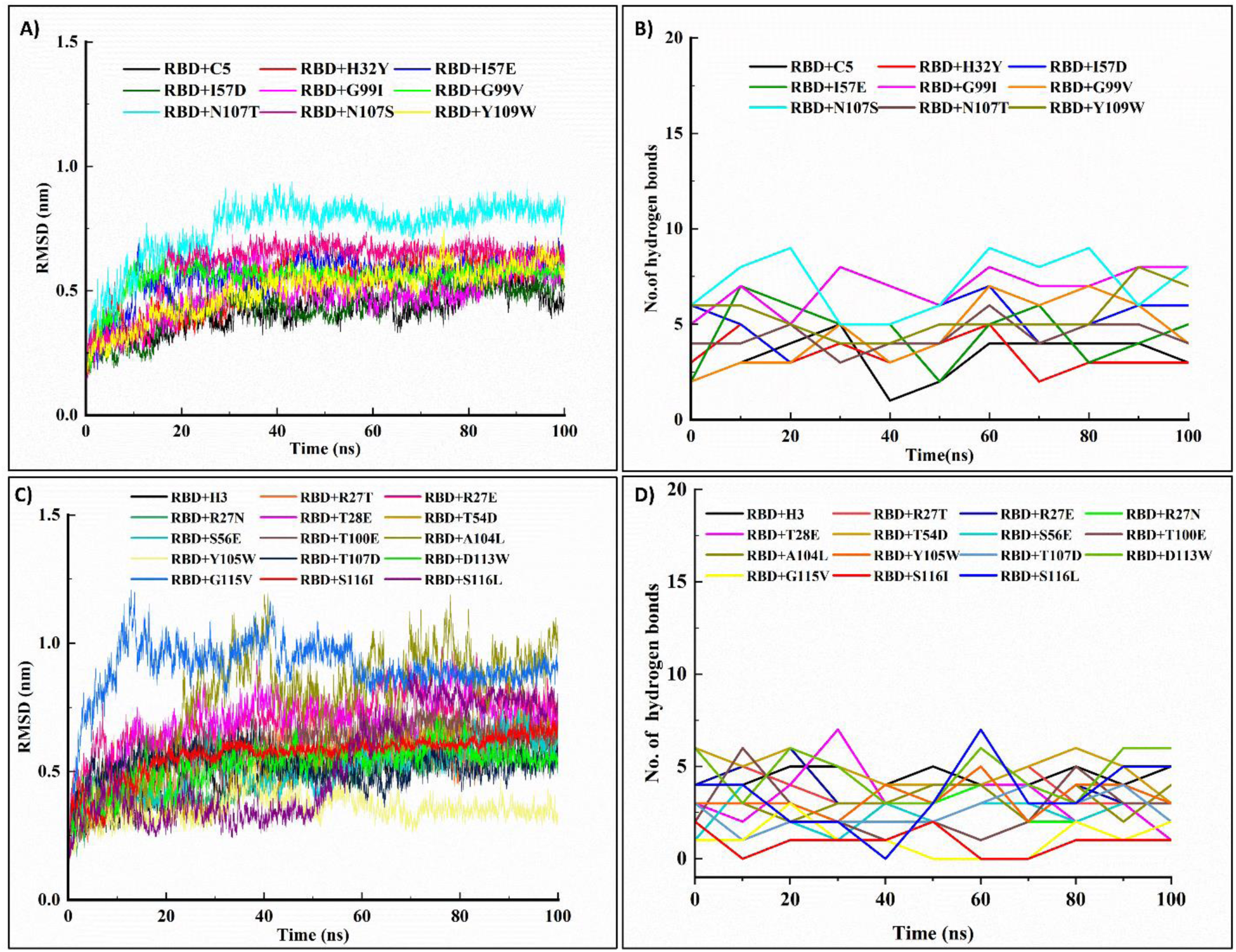
Molecular dynamics simulation analyses of the Omicron RBD complexed with nanobodies C5 and H3. RMSD value plots over 100 ns showing the stability of each RBD-nanobody complexes. Number of hydrogen bonds formed over time, indicating the interaction strength A-B) The plots of RMSD value and number of hydrogen bonds for C5 mutants in complex with Omicron RBD. C-D) The plots of RMSD value and number of hydrogen bonds for H3 mutants in complex with Omicron RBD.

**Supplementary Figure 3:**
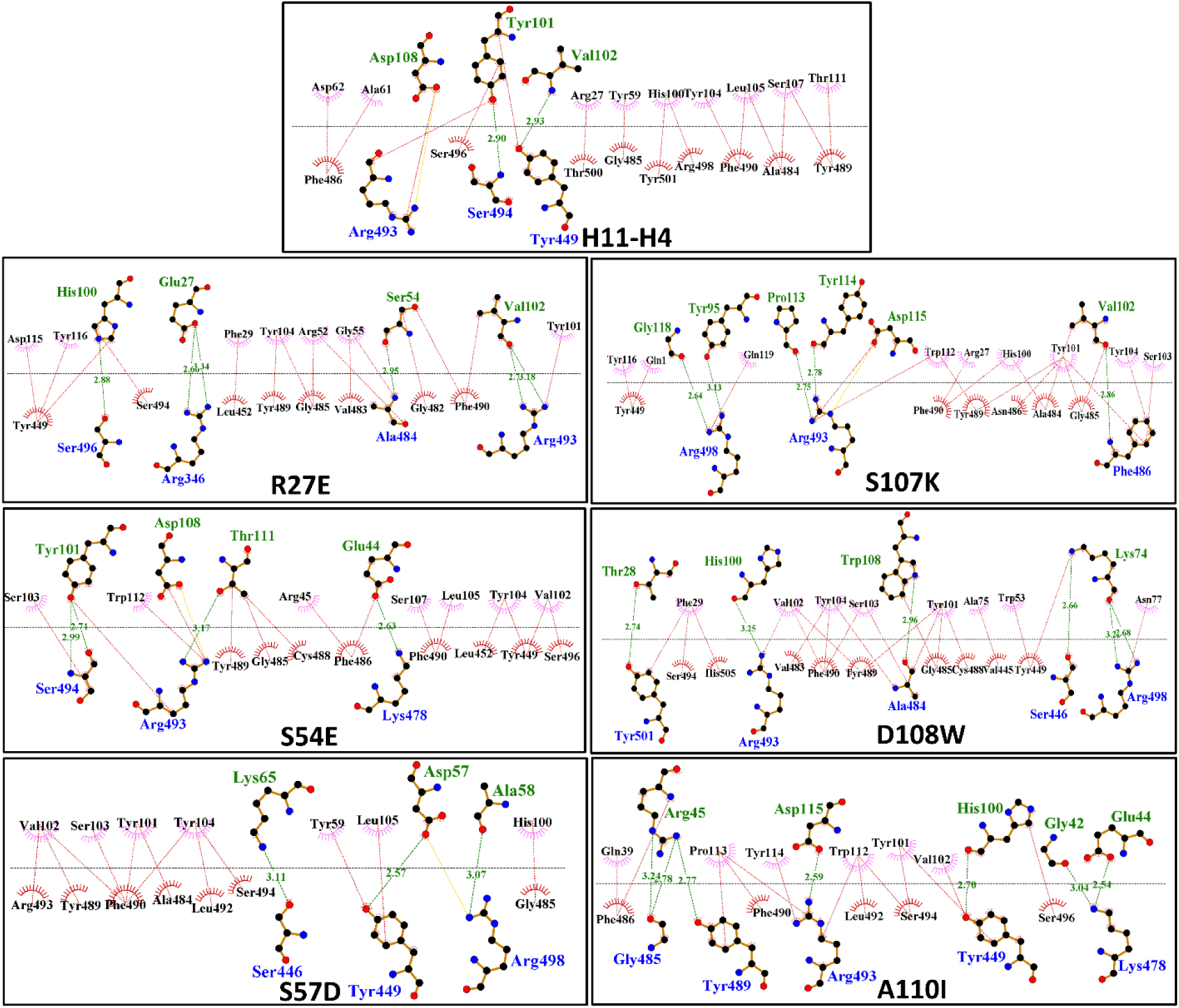
2D schematic representation of nanobody and RBD interacting residues generated through DimPlot. The docked complex of nanobody and RBD generated using HADDOCK 2.4 was used for the preparation of DimPlots. RBD residues were shown in green colour and nanobody residue were shown in blue colour. Hydrogen bonds are depicted by green dashed lines connecting the atoms involved, with the donor-acceptor distance indicated in Å. Salt bridges are depicted as yellow dashed lines and hydrophobic contacts are shown as red arcs with spokes extending towards the atoms they interact with.

**Supplementary Figure 4:**
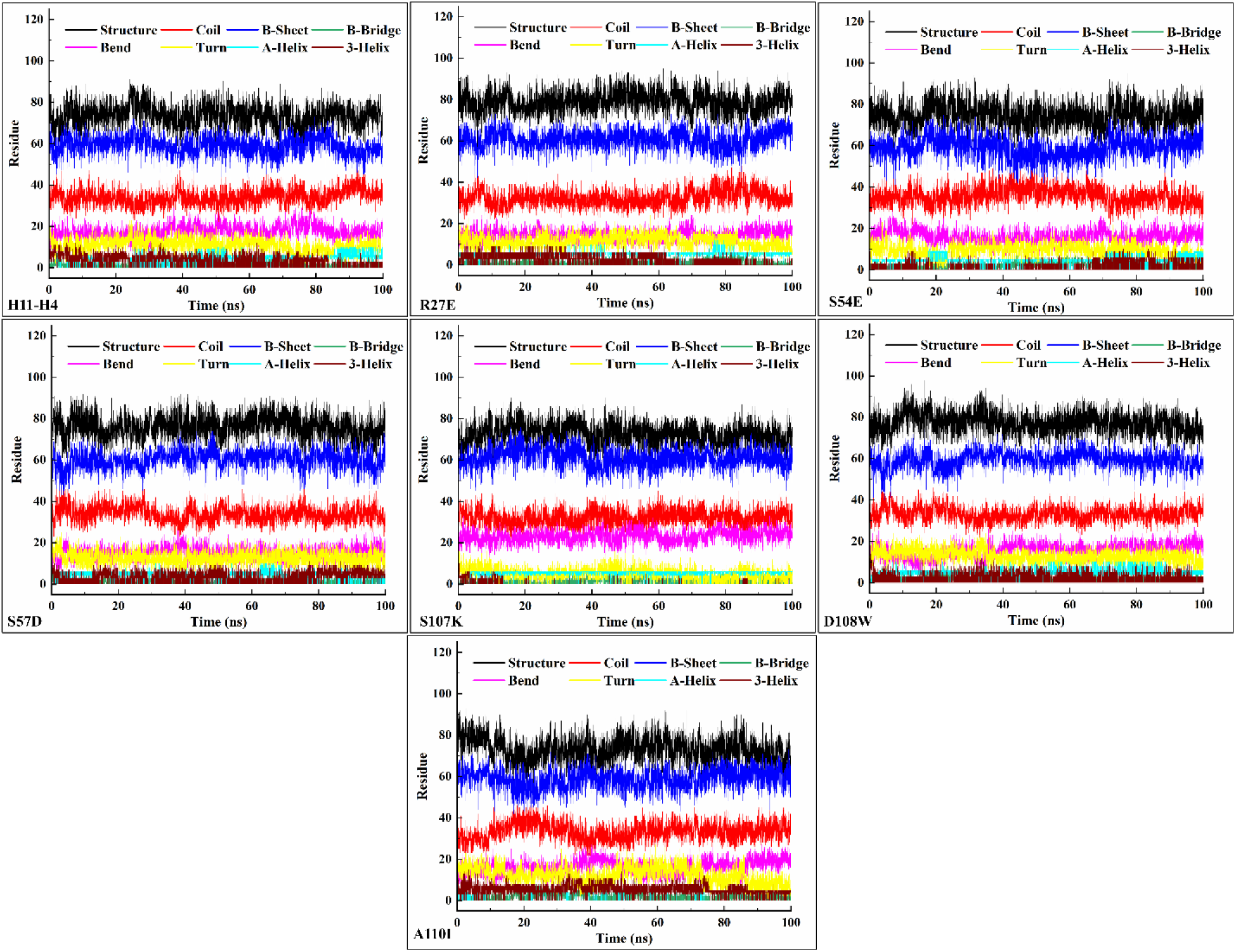
Secondary structure analysis of mutant nanobodies using gmx_dssp. Graphs of the relative number of residues in each secondary structure category plotted against time for the 100 ns MD simulation trajectory of nanobodies.

**Supplementary Figure 5:**
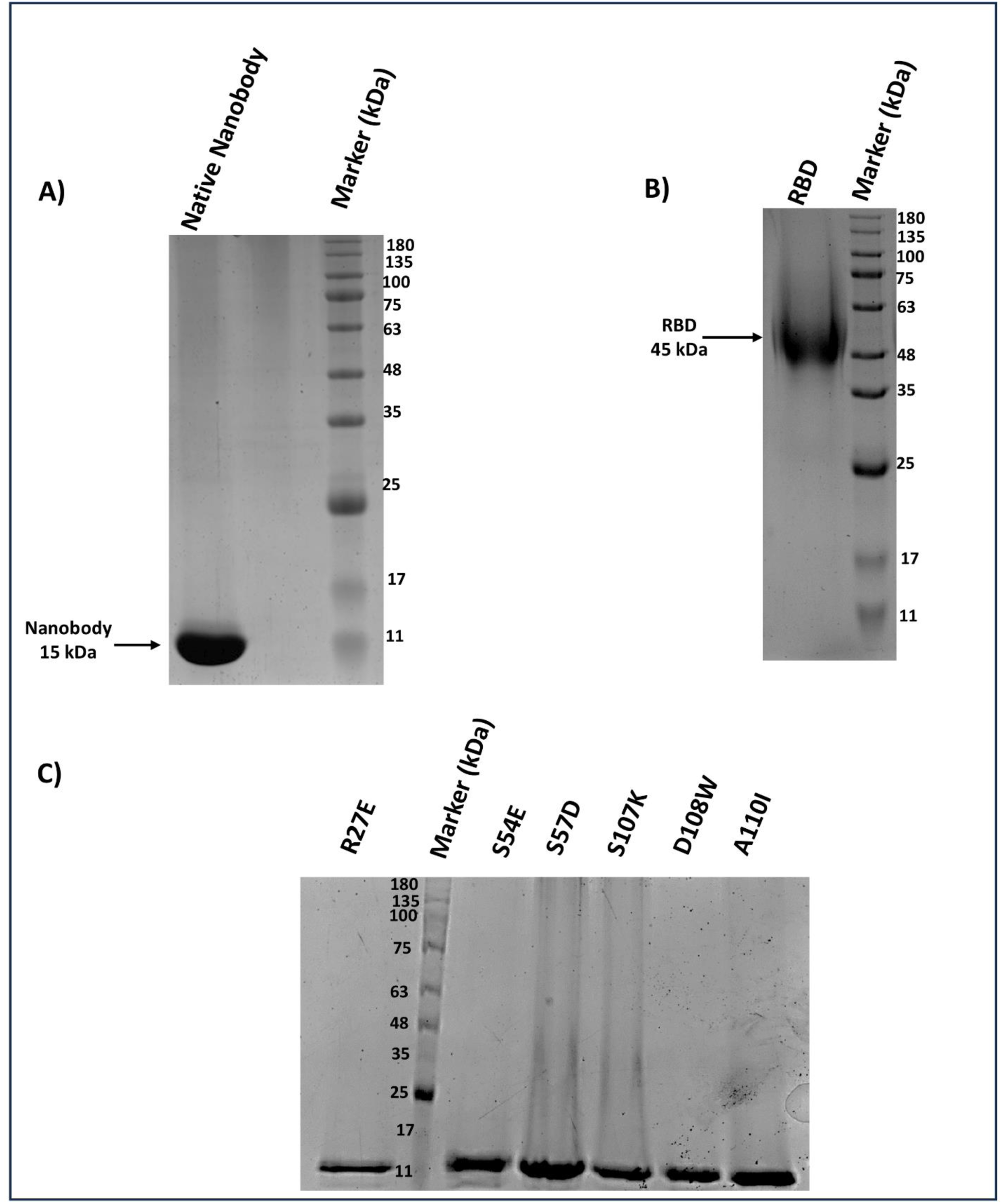
SDS-PAGE profile for the purification of H11-H4 nanobody, Omicron RBD and mutants of H11-H4 nanobody. A) Lane 1: marker (180 kDa); Lane 2: native nanobody (∼15 kDa). B) Lane 1: marker; Lane 2: RBD (∼ 45 kDa).C) Lane 1 H11-H4-R27E, Lane 2 Marker, Lane 3 H11-H4-S54E, Lane 3 H11-H4-S57D, Lane 4 H11-H4-S107K, Lane 5 H11-H4-D108W, Lane 6 H11-H4-A110I.

## Notes

### Competing Interest Statement

The authors have declared no competing interest.

